# Molecular Mechanisms Limiting the Therapeutic Window of AAV Gene Therapy in Mouse Models of Blue Cone Monochromacy

**DOI:** 10.1101/2025.02.14.638316

**Authors:** Brooke A. Brothers, Emily R. Sechrest, Li Ma, Madyson Ashcraft, Tongju Guan, Robert J. Barbera, Marion E. Cahill, Lee M. Shaw, Becky Chen, Wolfgang Baehr, Gangqing Hu, Peter Stoilov, Wen-Tao Deng

## Abstract

Blue cone monochromacy (BCM) is an X-linked retinal disorder caused by mutations in the *OPN1LW/OPN1MW* gene locus, resulting in impaired cone function and structural degeneration. We conducted a comparative analysis of AAV-mediated gene therapy in *Opn1lw/Opn1mw* double knockout (DKO) and *Opn1mw^C198R^/Opn1sw^-/-^* (C198R) BCM mouse models and evaluated the therapeutic window, efficacy, and longevity. Our results demonstrate that the AAV8-Y733F capsid achieved superior cone rescue compared to AAV5. While both DKO and C198R models showed similar therapeutic windows and rescue longevity, treatment efficacy decreased markedly in older mutant mice. Structural analysis revealed that aged cones in both models displayed degenerative changes, including mislocalized mitochondria and compromised connecting cilia. At the molecular level, we observed reduced AAV-mediated transgene expression in DKO and C198R older cones, which may result from decreased transduction efficiency, decreased circular episome stability, genome-wide transcription/translation downregulation, targeted mRNA/protein degradation, or overall cone degeneration. Notably, the cone-specific promoters for *Pde6c* and *Cngb3* maintained robust activity in degenerating cones. These findings suggest that combining an efficient AAV serotype with an optimized cone promoter could be a viable approach to extend the therapeutic window and enhance treatment longevity for BCM patients.

## INTRODUCTION

In the human retina, long (L-, red) and medium (M-, green) wavelength-sensitive cones comprise approximately 95% of the total cone population and are concentrated in the fovea, a small region in the retina responsible for high acuity vision in bright light conditions^1–3^. Their corresponding photo-sensitive proteins, L- and M-opsin, are encoded in the *OPN1LW/OPN1MW* gene locus on the X-chromosome^4^. Cone opsins are crucial for initiating the phototransduction cascade and for maintaining the structural integrity of cone outer segments (COS) where phototransduction occurs. Blue cone monochromacy (BCM) is an X-linked blinding disorder associated with mutations within the *OPN1LW/OPN1MW* gene locus and characterized by severely reduced or complete loss of function of red and green cone photoreceptors^5^. This disorder results in daytime blindness which involves loss of central vision, impaired color discrimination, and reduced visual acuity ranging from 20/60 to 20/200^6,7^.

One common class of mutation identified in BCM patients involves large deletions spanning the locus control region (LCR) of the *OPN1LW/MW* gene array and/or part of both genes, thereby preventing L/M-opsin expression. Previous studies demonstrate that BCM patients with deletion mutations exhibit significantly shortened COS as early as 5 years of age. However, mutant cones within these retinas appear to degenerate gradually throughout a patient’s life, evidenced by progressive foveal thinning tracked by optical coherence tomography (OCT) and Adaptive Optics Scanning Laser Ophthalmoscopy (AOSLO) longitudinally^6^. A second class involves various missense mutations in both *OPN1LW* and *OPN1MW* genes, resulting in translation of nonfunctional opsin proteins. The most common point mutation, predominantly identified in British populations, is a cysteine to arginine substitution at residue 203 (C203R) ^8–10^. Similar to deletion mutants, thinning of the fovea and shortening of COS are seen across the retinas of patients with C203R mutations (ages 5-70), but degeneration is less severe in early life compared to patients with deletion mutations^10^. Despite patients exhibiting significantly shortened COS, these studies show BCM patients retain intact cone inner segments (CIS) in the foveal region, which serve as potential targets for restoring normal cone opsin expression and rescuing cone function via gene therapy.

Previously, our lab generated mouse models that resemble either a BCM deletion (*Opn1mw^-/-^/Opn1sw^-/-^*, or DKO) or C203R (*Opn1mw^C198R^/Opn1sw^-/-^*, C198R) genotype. We performed AAV-mediated gene supplementation of human *OPN1LW* driven by a cone-specific PR2.1 promoter via subretinal injection. In both models, we observed robust rescue of cone function and regeneration of cone outer segment morphology when mice were treated at less than 3 months of age. However, therapy efficacy was significantly reduced when DKO and C198R mice were treated at 5 months or older^11,12^.

Previous studies have shown that success rates of AAV gene therapy for other diseases primarily affecting cones also exhibit age dependence. Restoration of cone-mediated vision using rAAV-based gene supplementation therapy has been reported in mouse, dog, sheep, and rhesus macaque models of *CNGA3*-, *CNGB3*-, or *PDE6C*-related achromatopsia. These studies demonstrated long-term restoration of cone-mediated vision^13–16^. However, decreases in therapy efficacy have been observed in aged mice, dogs, and rhesus macaques, underscoring the challenges of treating degenerating cones^13,14,16^. The potential causes have since been investigated, such as promoter silencing and histone modifications of the transgene^17–19^. The decreased rescue efficacy in aged photoreceptors is likely due to accumulated structural and molecular changes as a result of the mutation, leading to eventual photoreceptor death^20,21^.

In this study, we investigate the long-term efficacy of AAV gene therapy in DKO and C198R mice side-by-side and explore molecular mechanisms limiting the therapeutic window in aged cones. We demonstrate that therapeutic longevity and the treatment window are comparable between DKO and C198R mice. We detect severe morphological abnormalities in aging mutant cones and observe that aging cones exhibit significantly reduced transgene mRNA levels compared to younger cones. Additionally, we identify new candidate cone-specific promoters that may enhance transgene expression and improve therapeutic outcomes in older mice. Our findings offer insight into future studies that aim to expand the therapeutic window in translational BCM gene therapy research.

## RESULTS

### AAV8-Y733F-mediated *OPN1LW* expression improves rescue in C198R mice compared to AAV5

We have previously shown that DKO and C198R mice lack cone-mediated function, and that AAV5-mediated expression of *OPN1LW* driven by the cone-specific PR2.1 promoter successfully rescues cone function and COS structure^11,12^. To determine whether the AAV8-Y733F capsid, known for facilitating high photoreceptor transduction^22^, could mediate better rescue compared to AAV5, we treated C198R mice at 3 and 5 months of age with either AAV8-Y733F or AAV5 both expressing *OPN1LW* under the PR2.1 promoter. Electroretinograms (ERGs) were performed at 1 and 4 months post-injection in 3-month-old injected eyes (3M+1M and 3M+4M, respectively), and at 1 month post-injection in 5-month-old injected eyes (5M+1M).

AAV8-Y733F-treated C198R eyes demonstrated significantly better functional rescue compared to AAV5-treated eyes. Here, we show that 20% of AAV8-treated eyes had ERG amplitudes greater than 40 µV (N=11) in 3M+1M C198R animals, while only 5% of AAV5-treated eyes reached this level (N=10). In the 3M+4M group, 77.8% of AAV8-treated eyes showed ERG amplitudes above 40 µV (N=8), compared to 22.2% of AAV5-treated eyes (N=6). In contrast, only 12.5% of AAV8-treated eyes showed ERG amplitudes higher than 40 µV (N=10). In 5M+1M C198R animals, however, zero AAV5-treated eyes reached this level (N=12) (Fig. 1).

**Fig. 1.**
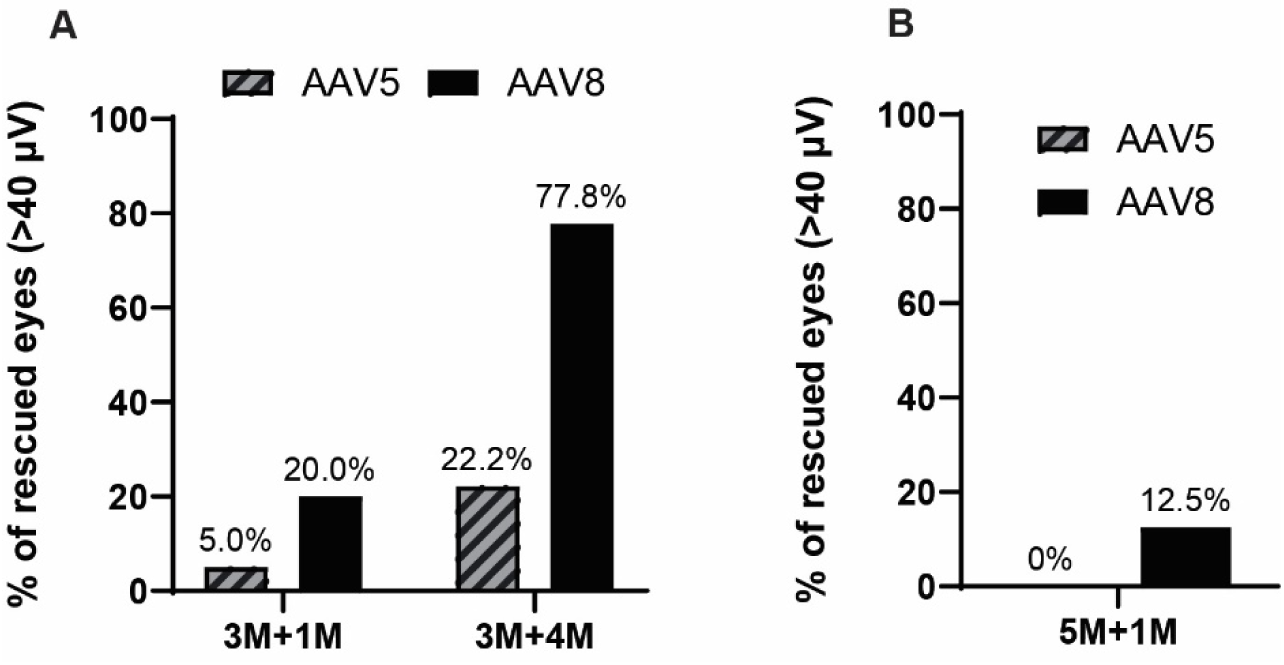
Comparison of rescue efficiency in C198R mice treated with AAV-mediated OPN1LW expression by serotypes AAV8-Y733F and AAV5. (A) C198R mice were treated at 3 months of age and assessed by ERG at 1- (3M+1M) and 4-months (3M+4M) post-injection. In 3M+1M, n=10 for AAV5, n=11 for AAV8-Y733F. In 3M+4M, n=6 for AAV5, n=8 for AAV8-Y733F. (B) C198R mice were treated at 5 months of age and assessed by ERG at 1- (5M+1M) post-injection. n=12 for AAV5, n=10 for AAV8-Y733F. In A and B, Y axis shows the percentage of treated eyes having b-wave maximum amplitudes higher than 40 µV.

### DKO and C198R mice exhibit comparable therapy rescue and therapeutic windows following AAV8-Y733F-mediated gene therapy

Next, we conducted a side-by-side study to directly compare the therapeutic efficacy, treatment window, and rescue longevity in DKO and C198R mice using the high-transduction AAV8-Y733F vector. Age-matched DKO and C198R mice were injected subretinally at 3, 5, and 7 months of age. Functional rescue was assessed by ERG at 1 month post-injection, with follow-up evaluations every 3 months until collection.

In both DKO and C198R mice treated at 3 months (Fig. 2A), we observed robust ERG rescue at 1 month post-injection (3M+1M). The average b-wave maximum amplitude at the highest tested light intensity of 25 cd·s/m^2^ was 23 μV ± 17 μV for DKO (n = 24), and 25 ± 16 μV for C198R (n=15). Functional rescue peaked at 4 months post-injection (3M+4M) with b-wave amplitudes of 55 ± 32 μV in DKO mice (n=11), and 47 ± 23 μV in C198R (n=10). At 7 and 10 months post-injection (3M+7M, 3M+10M), functional rescue persisted in both models, albeit with a slight decline compared to the 4-month time point. When ERG was assessed at 7 months post-treatment (3M+7M), average b-wave amplitudes were 23 ± 13 μV for DKO (n=10), and 42 ± 20 μV for C198R (n=10). At 10 months post-injection (3M+10M), average b-wave amplitudes were 13 ± 9 μV for DKO (n=5), and 35 ± 13 μV for C198R (n=7). Rescue efficiency appeared to be better maintained in C198R eyes compared to DKO eyes, though the difference was not statistically significant. The maximum amplitudes of ERG b-waves in treated DKO and C198R cones were significantly higher than those in untreated controls but remained significantly lower compared to wild-type (WT) controls (Fig. 2A; Fig. S1A).

**Fig. 2.**
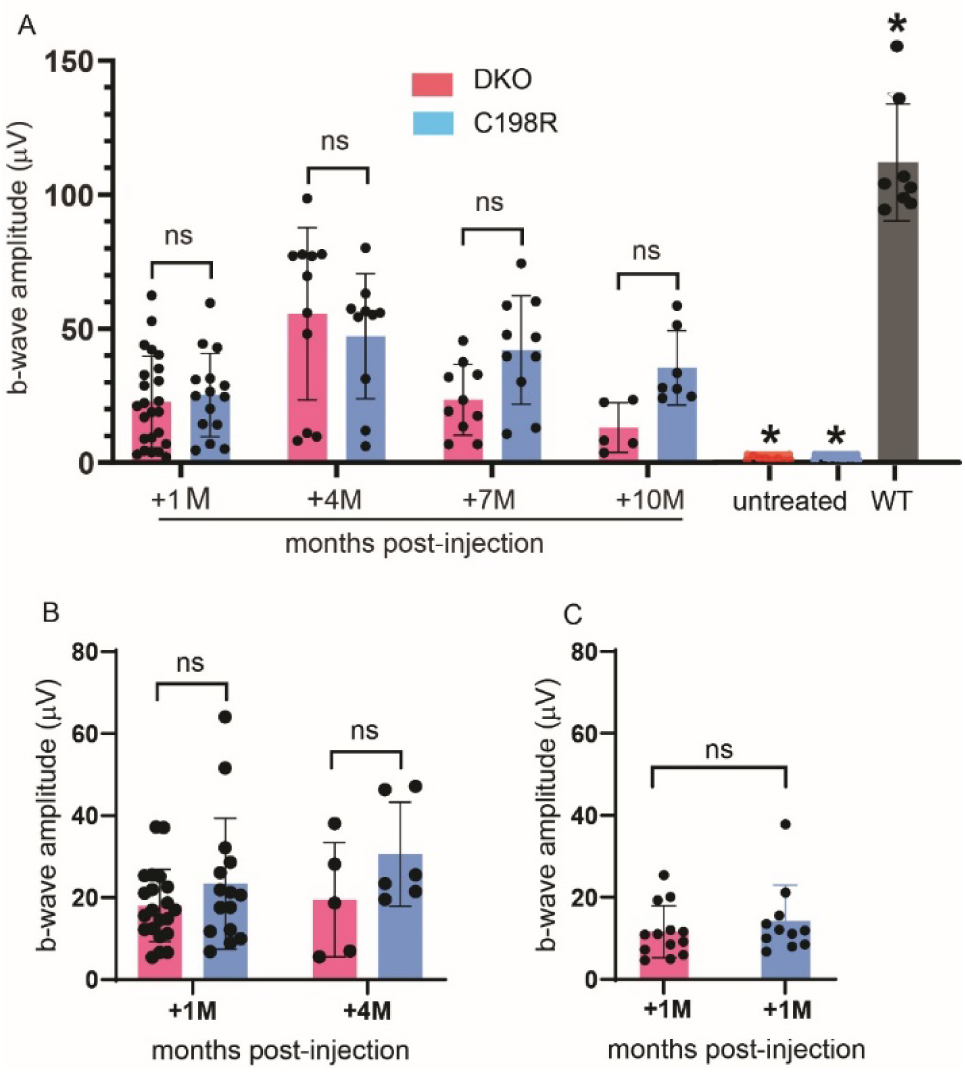
Cone-mediated b-wave maximum amplitudes from ERG responses of DKO and C198R mice treated at (A) 3 months, (B) 5 months, and (C) 7 months of age. (A) In 3M+1M mice, n=24 for DKO (pink bar), and n=15 for C198R (blue bar). In 3M+4M mice, n=11 for DKO, and n=10 for C198R. In 3M+7M mice, n=10 for DKO, and n=10 for C198R. In 3M+10M mice, n=5 for DKO, and n=7 for C198R. n=8 for WT, n=8 for DKO untreated, and n=8 for C198R untreated. All treated groups are significantly lower than WT (P < 0.05) but significantly higher than untreated controls (P < 0.05). Data shown are the average ± SD, 2-way ANOVA. (B) In 5M+1M mice, n=21 for DKO, and n=15 for C198R. In 5M+4M mice, n=5 for DKO and n=5 for C198R. There are no statistically significant differences between DKO and C198R mice in both time points. Data shown are average ± SD, 1-way ANOVA (P > 0.05). (C) In 7M+1M mice, n=13 for DKO and n=11 for C198R. There are no statistically significant differences between DKO and C198R mice. Data shown are average ± SD, non-paired t-test (P > 0.05).

In mice treated at 5 months of age, both DKO and C198R models showed similar levels of rescue at 1 and 4 months post-injection (5M+1M, 5M+4M; Fig. 2B, Fig. S1B). In 5M+1M mice, averaged b-wave amplitudes were 18 ± 9 μV and 23 ± 16 μV for DKO (n=21) and C198R (n=15) eyes, respectively. In 5M+4M mice, averaged b-wave amplitudes were 19 ± 14 μV for DKO (n=5) and 31 ± 13 μV for C198R (n=6) eyes. Rescue appeared to be maintained slightly better in 5M+4M C198R eyes, though the difference was not statistically significant.

Rescue efficiency of eyes treated at 7 months of age proved to be much lower than in those treated at 3 months, with b-wave amplitudes of 11 ± 6 μV and 14 ± 9 μV in DKO (n=13) and C198R (n=11) eyes, respectively (Fig. 2C, Fig. S1C). This is consistent with our previous findings that gene therapy is less effective in older mice^11,12^

Next, we performed immunohistochemistry (IHC) on treated eyes to confirm functional rescue was accompanied by structural rescue. COS morphology and cone phototransduction proteins were analyzed in retinal cross-sections from 3M+10M and 5M+1M treated DKO and C198R mice using antibodies against L/M-opsin, PDE6H (cone PDE γ-subunit), and GNAT2 (cone transducin α-subunit). In WT retinas, L/M-opsin, GNAT2, and PDE6H were abundantly expressed and localized primarily to COS (Fig. 3A and 3B, top row). In contrast, untreated DKO and C198R retinas showed no expression of M-opsin or GNAT2, and minimal PDE6H expression was detected but mislocalized to the CIS (Fig. 3A and 3B, second and third rows). In contrast, all treated eyes showed AAV-mediated L-opsin expression that was correctly localized to the COS (Fig. 3A and 3B, bottom four rows). Additionally, treatment partially restored PDE6H and GNAT2. Eyes treated at 3M+10M displayed more abundant expression of these three proteins compared to 5M+1M treated eyes, consistent with the observed increase of ERG b-wave amplitudes in 3-month-old treated mice.

**Fig 3.**
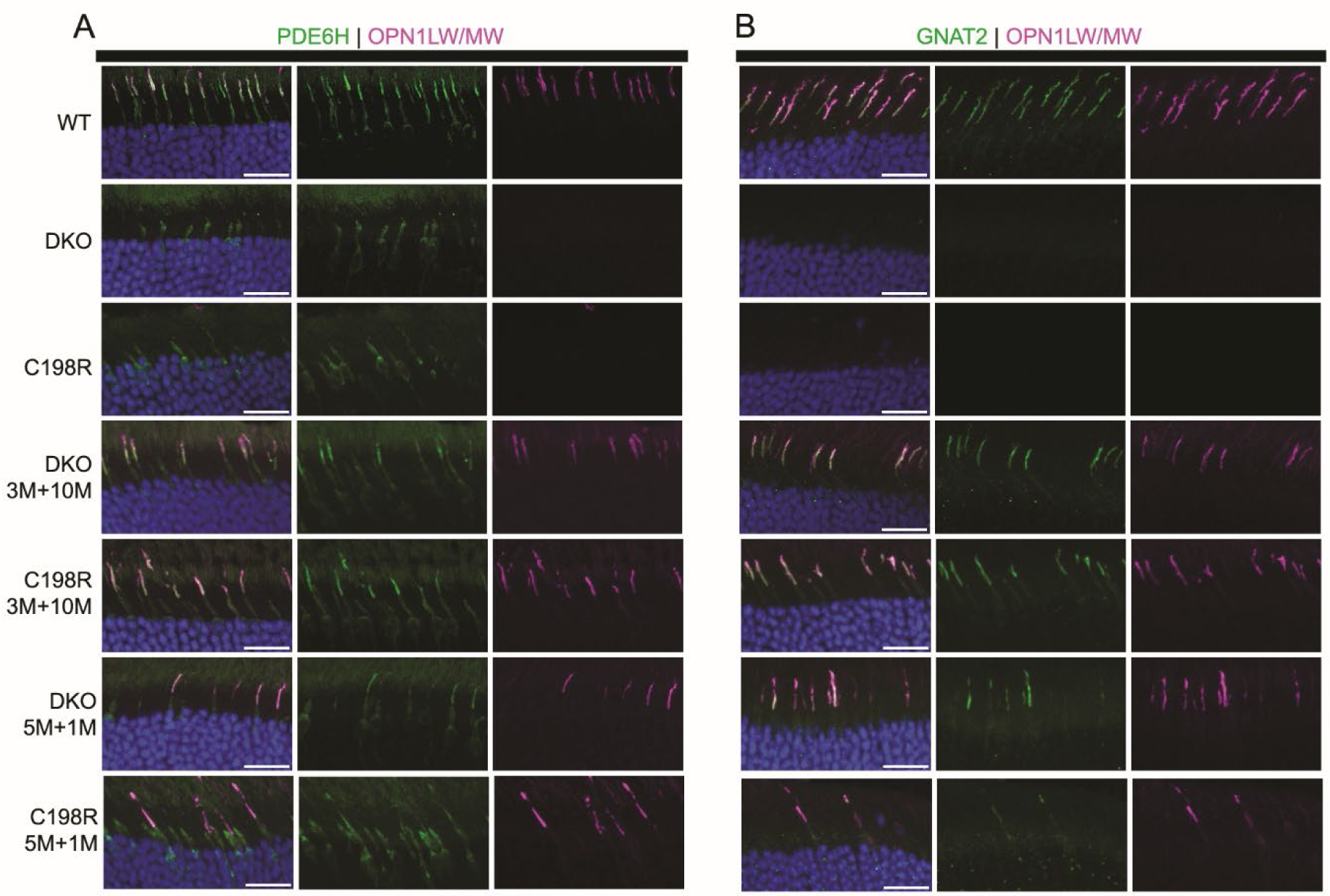
Immunohistochemistry analysis of retinal cross-sections stained with antibodies against OPN1LW/MW (magenta), PDE6H (green, A) and GNAT2 (green, B) from WT controls, untreated DKO, untreated C198R, and DKO and C198R eyes treated at 3- months of age and analyzed at 10 months post-injection (3M+10M), and treated at 5-months of age and analyzed at 1-month post-injection (5M+1M). Scale bar = 20 µm.

### Examination of cone ultrastructure before and after gene therapy

To closely examine cone ultrastructural changes before and after treatment, we performed transmission electron microscopy (TEM) on retinas from 1M+1M treated DKO and C198R mice, as well as untreated controls. Identifiable structures in WT cones included organized membrane discs in the COS, the connecting cilia (CC), the basal body (BB) which is localized between the COS and CIS at the base of the axoneme, as well as mitochondria concentrated directly below the CC in the CIS (Fig. 4A). In untreated 1-month-old DKO and C198R cones, COS were either absent or significantly shortened. A majority of cones at this age had intact CC and BB, along with mitochondria of typical size and concentration. However, some cones exhibited degenerating mitochondria that had begun migrating toward the middle and/or base of the CIS (Fig. 4B, 4D). In 1M+1M treated DKO and C198R cones, we observed varying degrees of COS regeneration — some cones exhibited well-organized membrane structures, while others remained disorganized and contained abnormal vacuoles. Additionally, both healthy and abnormal mitochondria were observed in the treated cones (Fig. 4C, 4E).

**Fig 4.**
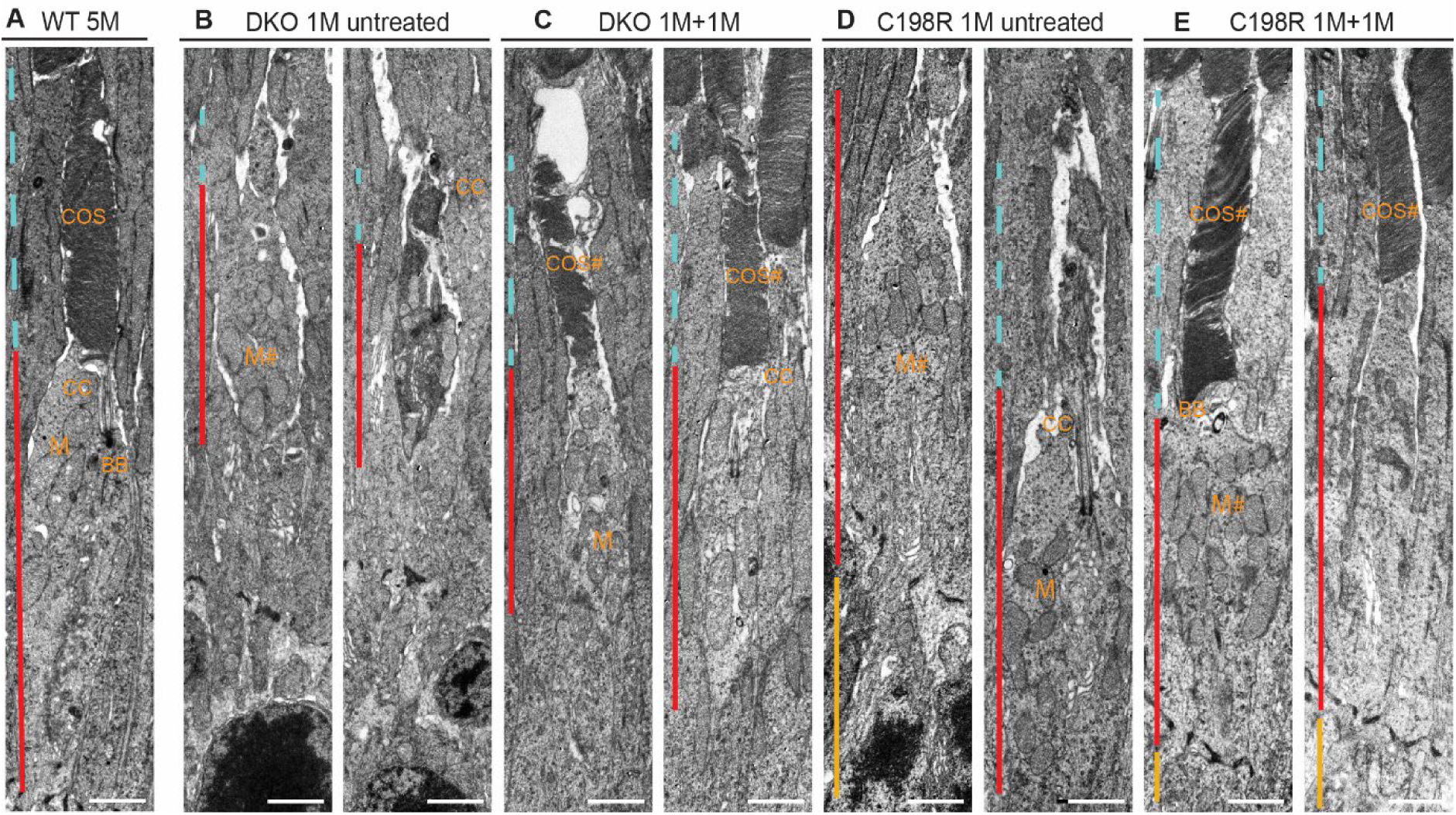
TEM images showing cone ultrastructure of WT, untreated DKO and C198R, and DKO and C198R treated at 1 month and analyzed at 1 month post-injection (1M+1M). (A) WT cone showing normal cone outer segment (COS, blue dashed line), cone inner segment (CIS, red line), basal body (BB), connecting cilium (CC), and M (mitochondria). (B, D) Untreated 1-month old DKO and C198R cones display disorganized, shortened, or missing COS (blue dashed line). Some with normal CIS, intact CC, and some with mild CIS (red line) degeneration with abnormal (M#) mitochondria. (C, E) DKO and C198R cones treated at 1-month of age and analyzed at 1-month post-injection demonstrate elaboration of COS (COS#), with varying phenotypes of normal (M) or abnormal (M#) mitochondria. Cell bodies/nuclei can be seen in some images (yellow line). Scale bar = 2μm

### Disease mechanisms underlying reduced therapy efficacy in aged mutant cones

We have previously shown and were able to reaffirm in the current study that gene therapy is less effective in aged DKO and C198R mice, likely due to progressive structural and molecular changes caused by the mutation^11,12^. To gain a better understanding of the ultrastructural changes in aged cones, we examined untreated DKO and C198R cones at 5 months of age using TEM, an age at which we previously observed significantly reduced therapeutic rescue^11,12^. While we identified a population of cones with intact CC and BB, we also observed cones with clear signs of degeneration. These degenerating cones exhibited mislocalized BB, located toward the middle or base of the CIS, as well as increased mislocalization of mitochondria that are normally concentrated at the apex of CIS but instead migrated to the middle and basal regions (Fig. 5).

**Fig. 5.**
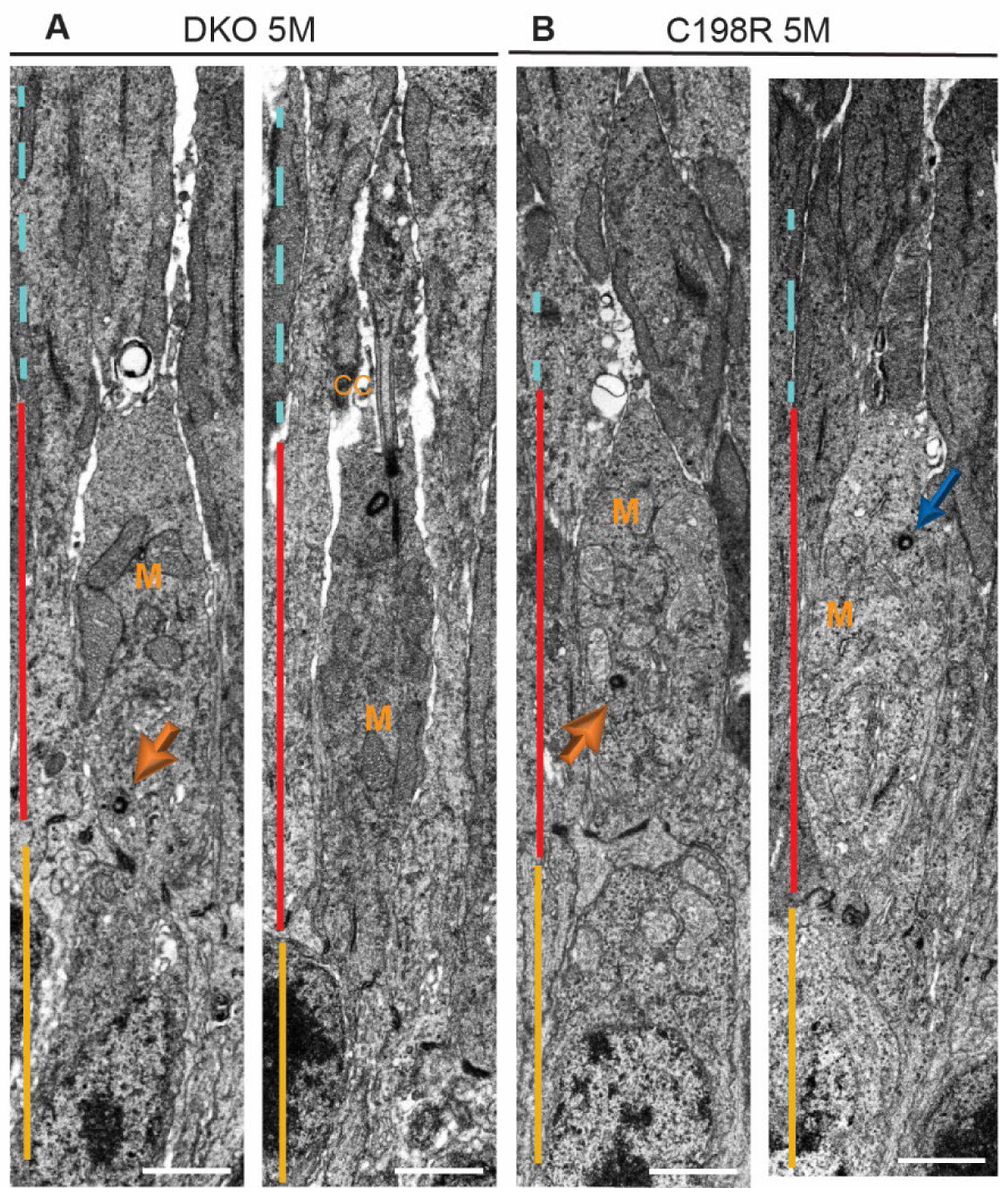
Representative TEM images of 5-month-old untreated DKO and C198R cones. (A) DKO and (B) C198R cones show more severe CIS defects many contain abnormal mitochondria (M#) and partially (blue arrow) or highly mislocalized BB (yellow arrow). Scale bar = 2μm.

We have also previously observed fewer cones expressed AAV-transduced L-opsin in DKO and C198R mice when treated at older ages^11,12^. To better quantify the percentage of cones expressing L-opsin and other COS proteins in eyes treated at younger versus older ages, we performed IHC on retinal flat mounts from 3M+1M and 7M+1M treated DKO and C198R mice, staining the cone sheath surrounding cones with peanut agglutinin (PNA), as well as L/M-opsin antibody that labels AAV-mediated L-opsin expression, and GNAT2 antibody that labels the cone transducin α-subunit (Fig. 6A; Fig. S2). We found that 3M+1M treated DKO and C198R retinas displayed similar numbers of viable PNA-positive cones, with 40% vs 48% of PNA+ cones expressing L-opsin in DKO vs C198R, respectively, and 28% vs 46% PNA+ cones expressing GNAT2 in DKO (n=3 mice, 7 areas counted) vs C198R (n=3 mice, 10 areas counted), respectively (Fig. 6B). Importantly, in 7M+1M treated eyes, viable cones expressing L-opsin and GNAT2 are much lower compared to 3M+1M cones. However, both DKO and C198R retinas again showed comparable numbers of viable cones and similar percentages of L-opsin^+^ and GNAT^+^ cones, with 6.5% vs 6.3% of PNA^+^ cones expressing L-opsin in DKO vs C198R treated eyes, respectively, and 12.2% vs 15.8% of PNA^+^ cones expressing GNAT2 in DKO (n=3 mice, 8 areas counted) vs C198R (n=3 mice, 8 areas counted) treated eyes, respectively (Fig. 6B).

**Fig. 6.**
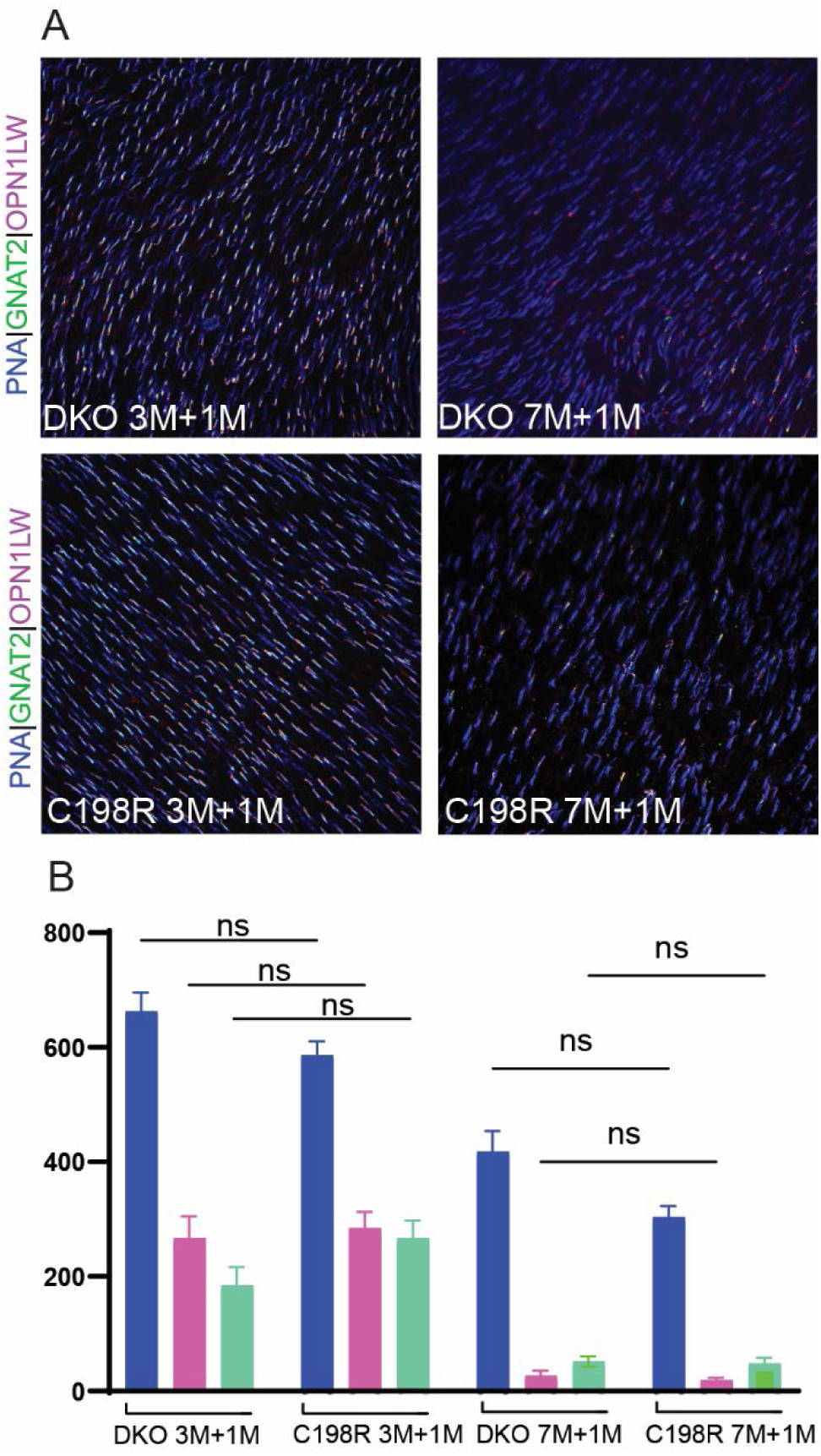
Comparison of AAV transduction efficiency of 3M+1M vs 7M+1M treated DKO and C198R cones by flat mounts. (A) Representative images of flat mounts stained with PNA (blue), antibodies against OPN1LW/MW (magenta), and GNAT2 (green). (B) Quantification of cone cells positive for PNA, OPN1LW, and GNAT2 in 3M+1M and 7M+1M DKO and C198R mice. N=3 retinas for each age and treatment group and 7-10 areas are counted. Unpaired t-test, P > 0.05.

To investigate if the significantly reduced L-opsin^+^ cones in mice treated at older ages are associated with decreased mRNA levels of AAV-mediated *OPN1LW* expression, we measured its mRNA levels in 1M+1M vs 4M+1M treated DKO and C198R eyes by qRT-PCR. The primers were designed to specifically amplify AAV-mediated human *OPN1LW,* rather than endogenous mouse *Opn1mw* (Fig. S3). The primer also did not produce any signal from wild-type mouse retina cDNA by qRT-PCR. qRT-PCR showed that *OPN1LW* mRNA levels in 4M+1M DKO and C198R mice were reduced to 45% (*P < 0.05, N=3) and 68% (P > 0.05, N=3-4) compared to 1M+1M DKO and C198R mice, respectively (Fig. 7). These results demonstrate that AAV-mediated *OPN1LW* mRNA expression is lower in mice treated at older ages.

**Fig. 7.**
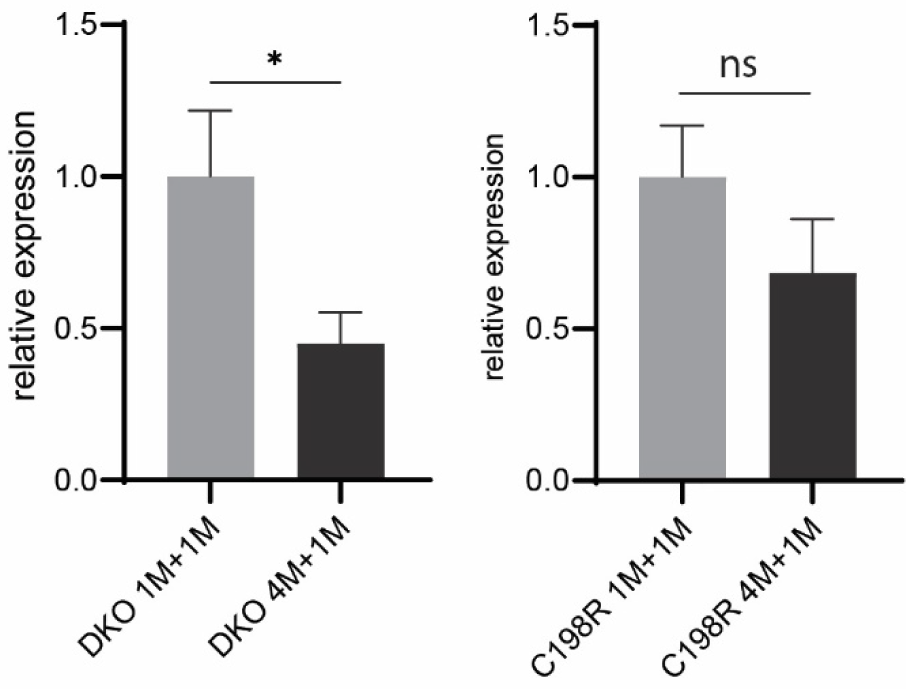
mRNA levels of AAV-mediated hOPN1LW in 1M+1M vs 4M+1M treated DKO (A) and C198R (B) by qRT-PCR. Data show average ± SEM. (A) N=3, *P < 0.05, non-paired t-test. (B) N=3-4, ns=not significant.

### Cone-specific promoters with robust activity in degenerating DKO and C198R cones

Another possible reason gene therapy is less effective in aged cones is that the PR2.1 promoter, derived from the promoter region of the human *OPN1LW/OPN1MW* gene locus^23^, may be less efficient in degenerating cones. To identify new potential candidate promoters for driving expression of the *OPN1LW* transgene in older mice, we investigated cone-specific genes whose expression remains stable or even increases in degenerating cones. We analyzed mRNA levels of seven cone phototransduction pathway genes that are abundantly expressed in normal COS, including *Opn1mw, Cngb3* (cone CNG β-subunit)*, Cnga3* (cone CNG α-subunit)*, Pde6c* (cone phosphodiesterase α-subunit)*, Pde6h* (cone phosphodiesterase γ-subunit)*, Gnat2, and Gngt2* (cone transducin γ-subunit), from single-cell RNA-sequencing (scRNAseq) data of 1- and 4-month-old DKO and C198R retinas. Among these genes, *Cngb3* and *Pde6c* maintained their expression levels in 4-month-old DKO and C198R cones compared to 1-month-old cones (Fig. 8A). To confirm these findings, we performed RT-qPCR in 1- and 4-month-old DKO, C198R, and WT retinas using *Pde6c* and *Cngb3* primers. RT-qPCR shows that 1- and 4-month-old DKO and C198R mice have similar mRNA levels for both *Pde6c* and *Cngb3* (P > 0.05). Moreover, they exhibit 3- to 5- and 2- to 3-fold higher mRNA levels compared to 1-month-old WT controls for *Pde6c* and *Cngb3* (*P < 0.01, N=3 samples/group, each sample is from 2 retinas of 2 different mice), respectively (Fig. 8B). These results suggest that *Pde6c* and *Cngb3* promoters may provide an alternative for robust transgene expression in mutant opsin-expressing cones at older ages.

**Fig 8.**
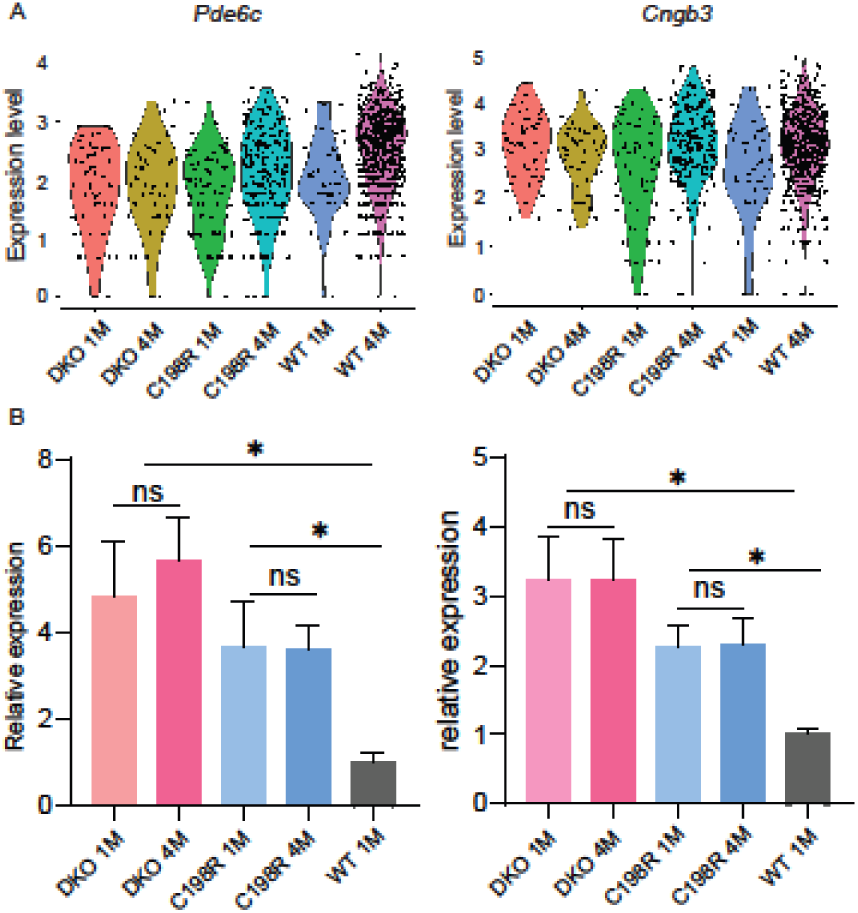
*Pde6c* and *Cngb3* maintain expression levels in 4-month-old DKO and C198R cones. (A) scRNA seq data showing *Pde6c* and *Cngb3* transcript levels in DKO 1- and 4-month-old, C198R 1- and 4-month-old, and WT 1- and 4-month-old retinas. (B) RT-qPCR showing mRNA levels of *Cngb3* and *Pde6c* in DKO 1- and 4-month-old, C198R 1- and 4-month-old, and WT 1-month-old retinas. N=3 per group, *P < 0.05, 1-way ANOVA.

## DISCUSSION

Our study presents a novel side-by-side comparison of AAV-mediated gene therapy in DKO and C198R mouse models, representing the two most prevalent causes of BCM. We evaluated therapeutic efficacy, window of treatment, and duration of effects, while also investigating the mechanisms that limit treatment effectiveness in aged cone cells. Our results reveal comparable therapeutic windows and treatment longevity between DKO and C198R mice. To improve therapeutic outcomes in aged cones, we showed that combining the enhanced transduction by the AAV8-Y733F capsid with a cone-specific promoter (*Pde6c* or *Cngb3*) could offer promising strategies for therapeutic advancement.

Decreased therapy efficacy in diseases primarily affecting cones has been observed in aged mice, dogs, and rhesus macaques^13,14,16^. Given the variable outcomes in clinical trials targeting cone diseases associated with *CNGB3*, *CNGA3*, *ABCA4* (ATP binding cassette subfamily A member 4), *CEP290* (centrosomal protein 290), and *GUCY2D* (guanylate cyclase 2D, retinal) ^24^, addressing the therapeutic window is critical to understanding these inconsistencies. Reduced therapy efficacy in aging cones may result from several factors: 1) reduced transduction efficiency due to fewer AAV particles entering cells, 2) impaired endosomal uptake and release of AAV particles, 3) decreased overall cellular transcription/translation caused by molecular changes in aged cones, 4) targeted degradation of transgene mRNA or protein, and 5) structural deterioration in aging cones.

Quantification of PNA^+^OPN1LW^+^ cones in mice treated at a young (< 3 months) vs older age (> 5 months) revealed that significantly fewer older cones successfully expressed the transgene compared to younger cones. Additionally, we showed that AAV-mediated *OPN1LW* mRNA is significantly lower in mice treated at older ages. These findings suggest that limitations in therapy efficacy could arise prior to protein translation. Possible contributing factors could include a reduced capacity of aged cones to be transduced by AAVs, less efficient endocytosis, increased endosomal degradation of AAVs, or reduced efficiency of the PR2.1 promoter. These mechanisms are not mutually exclusive.

Furthermore, we observed significant structural abnormalities in CIS in 5-month-old DKO and C198R cones, including degenerating connecting cilia, mislocalized basal bodies, and mitochondria migrating toward the base of the CIS. Since OPN1LW and other COS proteins are synthesized in the CIS and must be transported unidirectionally to the COS via the connecting cilium, structural deterioration of the cilium could disrupt this trafficking process, further contributing to a reduced therapy efficacy in aging cones.

Teasing out the mechanisms underlying reduced therapy efficacy in aged cones requires quantifying molecular and structural changes at the single-cell level, which presents significant technical challenges considering cones are only 3% of total photoreceptor cells in mouse retina. One of our current goals focuses on profiling the structural integrity of CIS and connecting cilia at various ages in DKO and C198R mice, categorizing these changes, and correlating them with transcriptomic alterations. Another ongoing effort of our lab involves determining whether AAV transducibility is reduced in aged cones as a potential strategy to address the low rescue efficacy in older mice. Additionally, our current method for evaluating therapy outcomes relies on full-field ERG, which measures a composite outcome by pooling the responses of all cones. However, this approach inherently lacks sensitivity. Instead, a visual-guided behavioral test offers a better option for detecting vision improvement in animals treated at middle or late stages, as the results are more clinically relevant.

Prior research in human patients demonstrated that individuals harboring the C203R missense mutation experience a slower rate of degeneration compared to those with deletion mutations^10^. This observation is puzzling, as *in vitro studies* have shown that the C203R mutation causes protein misfolding, retention in the endoplasmic reticulum, and potential toxicity to cone photoreceptors^25^. However, in C198R mice, we found that the mutant opsin was undetectable, indicating that cones can degrade this protein very efficiently^12^. Our side-by-side comparison of DKO and C198R mice revealed that C198R mice may have a slightly extended therapeutic window and therapy longevity, although these differences were not statistically significant. Considering the short lifespan of mice and the intrinsic variability of AAV-mediated gene therapy, proving better outcomes in C198R mice remains challenging. Nonetheless, our findings corroborate patient studies^6,10^, confirming that the C198R mutation does not cause more severe cone degeneration than deletion mutations, and treatment in C198R mice showed similar if not better outcomes than DKO mice. Additionally, both C198R and DKO cones exhibited similar abnormalities at older ages, including defects in COS discs, degenerating connecting cilia, and mislocalized centrioles and mitochondria. These findings suggest that both models experience comparable cellular deterioration over time.

In summary, we demonstrated that the AAV8-Y733F serotype proves superior in its transduction and gene delivery efficacy to cones compared to AAV5. In addition, DKO and C198R mice showed similar therapeutic windows and longevity, and treatment efficacy declined significantly in older mice of both genotypes likely due to cone degeneration and reduced transgene expression. Lastly, the cone-specific promoters for *Pde6c* and *Cngb3* demonstrated the potential to enhance therapy outcomes in aged cones due to their robust, upregulated expression compared to younger mice.

## MATERIALS AND METHODS

### Animals

*Opn1mw^-/-^Opn1sw^-/-^* (DKO) and *Opn1mw^C198R^Opn1sw^-/-^* (C198R) mice were described previously^11,12^. We observed no differences in phenotype between males and females for both strains, and both male and female mice were used in all experiments. Wild-type control mice are the C57BL/6J background. All experimental procedures involving animals in this study were approved and conducted in strict accordance with relevant guidelines and regulations by the Institutional Animal Care and Use Committee at West Virginia University, the ARVO Statement for the Use of Animals in Ophthalmic and Vision Research, and the National Institutes of Health.

### AAV vectors

The AAV construct used in this study contains a PR2.1 promoter that drives expression of human *OPN1LW* (PR2.1-OPN1LW)^23^. This construct was packaged in AAV serotypes 5 and 8-Y733F and purified according to previously published methods at the University of Florida Ocular Gene Therapy Core^26^.

### Subretinal injection

The viral vector for injection was prepared by adding fluorescein dye (0.1% final concentration) to AAV at a concentration of 1 x 10^10^ vector genomes per microliter. Preceding injection, mouse eyes were dilated using Tropi-Phen (Phenylephrine HCl 2.5%, Tropicamide 1%) ophthalmic solution (Pine Pharmaceuticals, Tonawanda, NY). Subsequently, Mice were then anesthetized by intramuscular (IM) injection using ketamine (80 mg/kg) and xylazine (10 mg/kg) in sterile phosphate-buffered saline (PBS). A 25-gauge needle was then used to create a small incision hole at the coronal edge. Next, a transcorneal subretinal injection was performed using a 33-guage blunt-end needle attached to a 5 μl Hamilton syringe containing 1 μl of the prepared AAV. The injection was administered into the subretinal space, and the injection bleb was visualized by the dispersion of fluorescence behind the retina. Immediately following injection, eyes were treated with Neomycin/Polymixin B Sulfates/Bacitracin Zinc ophthalmic ointment (Bausch & Lomb, Inc., Tampa, FL). Lastly, Antisedan (Orion Corporation, Espoo, Finland) was administered via intraperitoneal injection (IP) to reverse anesthesia.

### Electroretinography

All mouse eyes were dilated using Tropi-Phen drops (Pine Pharmaceuticals, Tonawanda, NY) prior to electroretinography (ERG) testing. Following eye dilation, animals were anesthetized with isoflurane (5% in 2.5% oxygen) for 3-5 minutes and placed onto a heated stage (37°C) with a nose cone supplying isoflurane (1.5% in 2.5% oxygen) throughout testing. Eyes were lubricated with GenTeal gel (0.3% Hypromellose), and silver wire electrodes were positioned above the corneal surface. A ground electrode was placed in the animals’ tails and a reference electrode was implanted subcutaneously between the ears. ERG recordings were performed using the UTAS Visual Diagnostic System, which included the BigShot Ganzfeld, UBA-4200 amplifier and interface, and EMWIN software (version 9.0.0, LKC Technologies, Gaithersburg, MD, USA). Following a 5-minute light adaption period, photopic (cone) responses were measured using a 30 cd/m^2^ white background light. Cone-mediated ERG responses were recorded at increasing light intensities (0.4, 0.7, 0.9, 1.4 log cd·s/m^2^) following stimulation with long- (630 nm) wavelength light.

### Retinal whole mount preparation and cone quantification

Mice were humanely euthanized directly before collections. The Change-a-Tip Deluxe Cautery tool (Braintree Scientific, Braintree, MA) was used to mark the dorsal position of the eye above the coronal line and immediately enucleated. Next, a 16-gauge needle was used to poke a hole along the edge of the cornea and the eye was placed in a 24-well plate containing 4% paraformaldehyde (PFA) in 1X PBS for a 60-minute incubation period at room temperature. After incubation, a radial cut was made at the dorsal position towards the optic nerve. The cornea was excised by cutting around the coronal line and the lens was removed. The neural retina was then dissected away from the eyecup and placed into a 94-well plate containing 4% PFA. The retinas were washed in 1X PBS for 30 seconds and were then blocked in 3% bovine serum albumin (BSA) with 0.3% Triton-X-100 in 1X PBS for 2 hours at room temperature. Next, retinal tissue was labeled with biotinylated peanut agglutinin (PNA) at a 1:500 dilution (Vector Laboratories, Burlingame, CA), L/M-opsin antibody at a 1:500 dilution (Kerafast (EJH006), and GNAT2 at a 1:500 dilution (Invitrogen, PA5-24553) diluted in 1% BSA in 1X PBS overnight at 4°C. The following day, retinas were washed three times for 15 minutes in 0.05% Tween-20 in 1X PBS and incubated overnight with Fluorescein Avidin D at a 1:500 dilution (Vector Laboratories, Burlingame, CA), Donkey anti-Rabbit IgG (H+L) Highly Cross-Adsorbed Secondary Antibody, Alexa Fluor™ 488 (Thermo Fisher, A-21206, 1:500) for GNAT2, and Goat anti-Chicken IgY (H+L) Cross-Adsorbed Secondary Antibody, Alexa Fluor™ Plus 594 (Thermo Fisher, A32759) for L/M opsin. Finally, retinas were washed, and three additional radial cuts were made at the ventral, temporal, and nasal edges of the retina. The retinal whole mount was then flattened under a coverslip and mounted using ProLong Gold Antifade Mountant (Thermo Fisher, Waltham, MA). Whole mounts were imaged using a Nikon C2 confocal microscope and processed using ImageJ FIJI software. Cells positive for PNA, GNAT2, and L/M-opsin were counted using ImageJ software. Three retinas from each group were processed, imaged, and counted.

### Immunohistochemistry and frozen retinal cross-section preparation

Mice were humanely euthanized directly before collections. The Change-a-Tip Deluxe Cautery tool (Braintree Scientific, Braintree, MA) was used to mark the dorsal position above the coronal line and immediately enucleated. Next, a 16-gauge needle was used to poke a hole along the edge of the cornea and the eye was incubated in 4% paraformaldehyde (PFA) in 1X PBS for 2 hours at room temperature. The cornea was then removed by cutting around the coronal line. Next, the eyes were washed three times in 1X PBS for 10 minutes and placed in 20% sucrose in 1X PBS overnight at 4°C. The next day, the eyes were incubated in a 50/50 mixture of 20% sucrose in 1X PBS and Tissue-Tek O.C.T compound (Sakura Finetek USA, Inc., Torrance, CA) for 1 hour at 4°C. After incubation, the eyes were positioned in a cryomold filled with Tissue-Tek® O.C.T. and flash frozen in a dry ice and ethanol bath. Before staining the retinal tissue, 16µm cross-sections were cut using a Leica CM1850 Cryostat and placed on Superfrost plus slides (Fisher Scientific). A hydrophobic PAP pen was used to draw a barrier around the tissue on the slides and then washed with 1X PBS to eliminate O.C.T. compound and to hydrate the retinal cross-sections. Tissue sections were then incubated in a blocking buffer containing 3% bovine serum albumin (BSA) and 0.3% Triton-X-100 for 1 hour at room temperature. Then, primary antibodies were diluted in 1% BSA in 1X PBS and incubated overnight at 4°C. The following day, cross-sections were washed three times for 7 minutes with 0.1% Triton-X-100 in 1X PBS, followed by one 7-minute wash with 1X PBS. After washing, secondary antibodies and DAPI (1:1000; Thermo Fisher, Waltham, MA) diluted in 1X PBS were added and incubated for 2 hours at room temperature. Following additional washes in 0.1% Triton-X-100 in 1X PBS, coverslips were mounted using ProLong Gold Antifade Mountant (Thermo Fisher, Waltham, MA). Retinal cross-sections were imaged using a Nikon C2 confocal microscope and processed using ImageJ FIJI software. Primary antibodies used are: PDE6H (Proteintech, Cat No. 18151-1-AP, 1:500 dilution), OPN1L/M (Kerafast, EJH006, 1:1000 dilution), and GNAT2 (1:500 dilution (Invitrogen, PA5-24553, 1:1000 dilution). Secondary antibodies used are Goat anti-Chicken IgY (H+L) Cross-Adsorbed Secondary Antibody, Alexa Fluor™ Plus 594 (Thermo Fisher, A32759, 1:500 dilution), Donkey anti-Rabbit IgG (H+L) Highly Cross-Adsorbed Secondary Antibody, Alexa Fluor™ 488 (Thermo Fisher, A-21206, 1:500 dilution).

### Ultrastructural analysis of cone photoreceptors

Retinal samples were prepared and imaged using previously published procedures^27–29^. Briefly, the enucleated eyes were fixed in a solution containing 2% paraformaldehyde and 2.5% glutaraldehyde in 100 mM cacodylate buffer (pH 7.4). A small incision was made at the edge of the cornea using a 20 G needle. The eyes were then incubated in a glass vial containing the fixative for 30–60 minutes at room temperature. Subsequently, the eyes were transferred to a petri dish with a drop of 7% sucrose in 200 mM cacodylate buffer (pH 7.4). After removing the cornea and lens, the eyecups were placed back into the fixative-filled glass vial and incubated for two days.

After fixation, the eyecups were sectioned into smaller trapezoid-shaped pieces and treated with 2% osmium tetroxide in 0.1 M cacodylate buffer, followed by incubation with 1% uranyl acetate. The fixed tissue was then dehydrated through a graded ethanol series and embedded in Polybed 812 resin (Polysciences, Inc.). Thin sections were mounted on grids, post-stained with 3% Reynold’s lead citrate, and imaged using a JEOL 1010 transmission electron microscope at 80 kV.

### RNA extraction, cDNA synthesis, and quantitative RT-PCR

Total RNA was extracted from mouse retinas using the Quick-RNA MicroPrep system (Zymo Research, R1050) according to the manufacturer’s protocol. Concentration and purification quality were measured by the Nanodrop ND-1000 Spectrophotometer. First-strand cDNA was synthesized using the iScript cDNA Synthesis kit (Bio-Rad, 1708891), and qRT-PCR was performed with iQ SYBR Green Supermix (Bio-Rad, 1708882), loading approximately 120 ng cDNA per reaction. 40 PCR cycles were run with an annealing temperature of 57°C. Fold change was calculated using the 2-ΔΔCT method. Actin as an endogenous control for all experiments.

Primers used to amplify OPN1LW transcripts in treated mice are: CTGCATCATCCCACTCGCT (forward) and GCTTTGCCACCGCTCG (reverse). ΔΔCT values were generated by normalizing 4M+1M treated replicates to 1M+1M treated. To amplify Pde6c transcripts, primer sequences were GGATAGTTGGCTGGGTCGCT (forward) and TGCCATGACCACAGCAAGGA (reverse). Cngb3 primer sequences were GCCAACCATAGCACAGGGAG (forward) and TGCTCGACATTCAGGGTCAG (reverse). Actin primers were used as internal controls and data normalization. Actin primers are: ACCAACTGGGACGACATGGAGAA (forward) and CATGGCTGGGGTGTTGAAGGT (reverse); Both sets of ΔΔCT values were generated by normalizing all DKO and C198R replicates to wild-type controls.

### Transcriptomics analysis

Single-cell suspensions were prepared with pooled retinas from one- and four-month-old male and female from each genotype according to the protocol by Fadl and colleagues to preserve cone viability^30^. Single-cell libraries were prepared using the 10x Chromium Platform following the manufacturer’s instructions at the WVU Flow Cytometry & Single Cell Core Facility. Sequencing was performed at the Admera Health (South Plainfiled, NJ). scRNA-seq FASTQ reads were aligned to the GRCm38 (mm10) reference genome and cellranger count (version 6.1.2) was used to perform alignment, filtering, barcode counting, and UMI (Unique Molecular Identifier) counting. Seurat (version 4.2.1) was used for downstream analysis^31^. All sample data are combined into a merged Seurat object. Normalize Data was used for normalization and set normalization method as ‘CLR’ (centered log ratio transformation). The cell cycle phase score was calculated by CellCycleScoring. FindVariableFeatures was used to calculate a subset of features that exihibit high cell-to-cell variation in the dataset. Focusing on these genes in downstream analysis helps to highlight biological signal in single-cell datasets. The top 2000 features were selected. Downstream principal component analysis (PCA) was then performed on the scaled data, use the determined variable features as the input. Data were scaled and centered by ScaleData function and regress out the cell cycle’s effect. RunPCA was used for principal component analysis (PCA) dimensionality reduction. RunUMAP was used for dimensional reduction and visualization via uniform manifold approximation and projection (UMAP). FindNeighbours was used to compute the nearest neighbors for the object. FindClusters was used to identify clusters of cells by a shared nearest neighbors modularity optimization based on original Louvain clustering algorithms. Each subclusters features were plotted to confirm quality. After filter out the subclusters which have a significant low level of RNA fragments, further quality control was based on each library’s features with these settings: 500 < nCount_RNA (total number of RNA fragments) < 20000, 500 < nFeature_RNA (number of genes) < 6000, percent.mt < 30%, percent.ribo < 15%. Percent.mt (RNA reads from mitochondrial genes) and percent.ribo (reads from ribosomal genes) were calculated by the PercentageFeatureSet function. After quality control, subclusters were identified based on the same subclusters identification procedures. Cone photoreceptor cells were annotated through marker genes, including Arr3, Cnga3, Cngb3, Gnat2, Opn1mw, and Pde6c. Then this specific population was extracted for analysis (Table S1). All raw sequenced data are publicly available at Gene Expression Omnibus under accession number GSE289547. Secure token for data access during manuscript peer-review is afifgeyyvbwbrub.

### Statistical analysis

All data are presented as the mean ± SD unless otherwise noted, and figure legends contain details about sample size. All analyses were carried out using GraphPad Prism 9 software by unpaired, two-tailed Welch’s t-test (2 groups), ordinary 1-way ANOVA with Turkey ad hoc test (more than two groups), or by two-way ANOVA with Turkey ad hoc test for multiple comparisons, unless noted differently. Significance is indicated as \**p* ≤ 0.05, ns = not significant.

## ACKNOWLEDGEMENTS

This work was supported by National Institutes of Health (R01 EY030056 to W.T. Deng, R01 EY08123, EY014800-039003 to W. Baehr, and R01 EY025536, R21 EY036144 to P. Stoilov), West Virginia University startup fund, NIH NIGMS P20GM144230 Visual Sciences COBRE grant to WVU and an unrestricted challenge grant from Research To Prevent Blindness (RPB) to the Ophthalmology department to WVU, BCM Families Foundation, unrestricted grants to the University of Utah Department of Ophthalmology from Research to Prevent Blindness (RPB), and the West Virginia Lions and Lions Club International Foundation. Data analysis support through the WVU Bioinformatics Core is supported by NIGMS P20 GM103434 and NIGMS U54 GM104942 (to G. Hu).

