## Supplemental for "Molecular Mechanisms Limiting the Therapeutic Window of AAV Gene Therapy in Mouse Models of Blue Cone Monochromacy"

### Supplemental Materials

Fig. S1

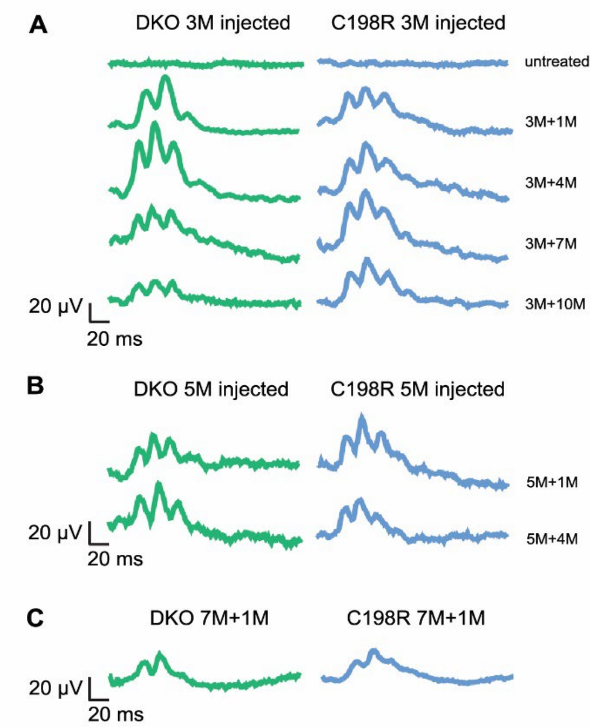


Fig S1. Representative ERG traces from DKO and C198R mice treated at different ages. Representative ERG waveforms of DKO and C198R mice treated at (A) 3 months, (B) 5 months, and (C) 7 months of age and assessed by ERG at 1, 4, 7, and 10 months post-injection.

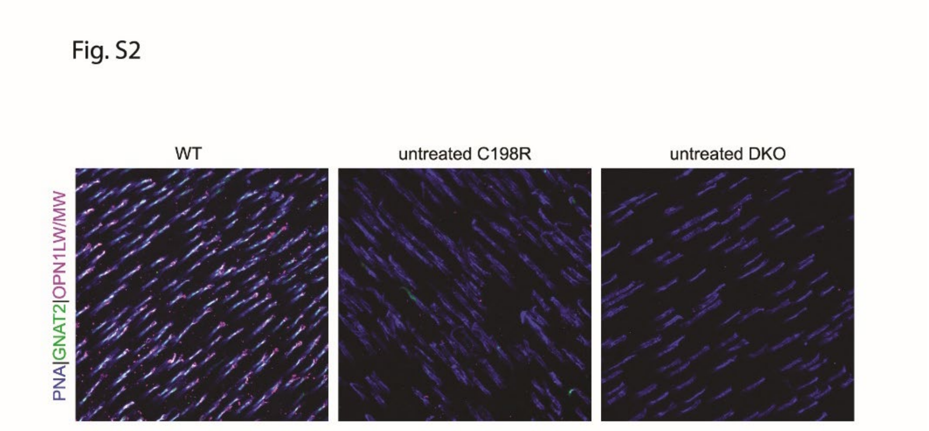


Fig. S2. Representative flat-mount images of WT, and untreated DKO and C198R retinas stained with PNA (peanut agglutinin, blue), antibodies against GNAT2 (green) and OPN1LW/MW (magenta).

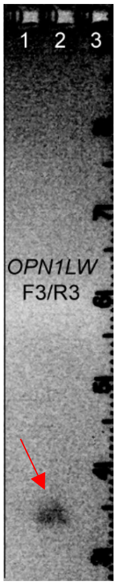


Fig. S3. OPN1LW primers are specific to human *OPN1LW* (arrow) not mouse *Opn1mw* cDNA. Agarose gel analysis of PCR product amplified using cDNA from wild-type mice (lane 1), and cDNA from DKO mice injected with AAV-PR2.1-hOPN1LW (lane 2), and no template control (lane 3) .

| barcode | sample |
|---|---|
| AGTTAGCGTTCCCACT-1 | C198R_1M_rep1 |
| ATGCGATGTCGGTGAA-1 | C198R_1M_rep1 |
| CAACCTCAGCATTGTC-1 | C198R_1M_rep1 |
| CACTTCGTCGCTCATC-1 | C198R_1M_rep1 |
| CAGCGTGTCACGGGCT-1 | C198R_1M_rep1 |
| CATCGTCTCGGAGTGA-1 | C198R_1M_rep1 |
| CCACACTAGAATCCCT-1 | C198R_1M_rep1 |
| CTCCTCCAGAAATCCA-1 | C198R_1M_rep1 |
| CTTACCGCAGCAGTCC-1 | C198R_1M_rep1 |
| GCATTAGAGCCGTTAT-1 | C198R_1M_rep1 |
| GCGATCGGTAGACAGC-1 | C198R_1M_rep1 |
| TACGGGCAGGAACATT-1 | C198R_1M_rep1 |
| TCCTCTTCAAGTTGGG-1 | C198R_1M_rep1 |
| TGTTGGAGTGAACGGT-1 | C198R_1M_rep1 |
| TTAGGGTGTATCCCTC-1 | C198R_1M_rep1 |
| AAGCATCGTAGATTAG-1 | C198R_1M_rep2 |
| ACCTGTCCAAGCAGGT-1 | C198R_1M_rep2 |
| AGAGAATAGTACAGAT-1 | C198R_1M_rep2 |
| AGCCAATCATCGATCA-1 | C198R_1M_rep2 |
| AGGACGATCCCATTCG-1 | C198R_1M_rep2 |
| AGGACTTTCGTGCTCT-1 | C198R_1M_rep2 |
| AGGTGTTGTCATCCCT-1 | C198R_1M_rep2 |
| AGGTGTTTCATGCCGG-1 | C198R_1M_rep2 |
| CAACGATAGATAACGT-1 | C198R_1M_rep2 |
| CACTAAGCAAATGCTC-1 | C198R_1M_rep2 |
| CACTTCGTCGTGCGAC-1 | C198R_1M_rep2 |
| CAGGCCATCGCACGGT-1 | C198R_1M_rep2 |
| CATCGGGAGGAAGTCC-1 | C198R_1M_rep2 |
| CATTCCGTCCATCACC-1 | C198R_1M_rep2 |
| CCACTTGTCGCTATTT-1 | C198R_1M_rep2 |
| CCTGTTGAGATGCAGC-1 | C198R_1M_rep2 |
| CGATCGGAGCGAGTCA-1 | C198R_1M_rep2 |
| CGGCAGTGTATTTCTC-1 | C198R_1M_rep2 |
| CGGCAGTTCATGAGGG-1 | C198R_1M_rep2 |
| CTCAATTAGATCCAAA-1 | C198R_1M_rep2 |
| CTCATTACACGCGCAT-1 | C198R_1M_rep2 |
| CTGAGGCAGTCACTGT-1 | C198R_1M_rep2 |
| CTTAGGACATATCTCT-1 | C198R_1M_rep2 |
| CTTCCTTAGCACTAGG-1 | C198R_1M_rep2 |
| GAAACCTAGCTCACTA-1 | C198R_1M_rep2 |
| GACTGATAGGAGGGTG-1 | C198R_1M_rep2 |

| | |
|---|---|
| GACTGATCAACGTATC-1 | C198R_1M_rep2 |
| GAGGCAAGTGGTCTTA-1 | C198R_1M_rep2 |
| GAGTGTTTCTGTCTCG-1 | C198R_1M_rep2 |
| GAGTTGTAGAATTGCA-1 | C198R_1M_rep2 |
| GCGATCGGTCGCTTAA-1 | C198R_1M_rep2 |
| GTAGTACTCAGCAATC-1 | C198R_1M_rep2 |
| GTGCACGTCTCGTCAC-1 | C198R_1M_rep2 |
| GTGTGGCCAGCGGTCT-1 | C198R_1M_rep2 |
| TAACCAGAGGAGAATG-1 | C198R_1M_rep2 |
| TAATTCCTCCATGCAA-1 | C198R_1M_rep2 |
| TATTGGGGTCGTTGCG-1 | C198R_1M_rep2 |
| TATTTCGAGTGCGACA-1 | C198R_1M_rep2 |
| TCACAAGGTATGGTAA-1 | C198R_1M_rep2 |
| TCACACCGTCCATAGT-1 | C198R_1M_rep2 |
| TCACGCTGTATCTTCT-1 | C198R_1M_rep2 |
| TCAGGGCAGTTGCTGT-1 | C198R_1M_rep2 |
| TCATGTTGTCACAGTT-1 | C198R_1M_rep2 |
| TCCACCAGTCACTCGG-1 | C198R_1M_rep2 |
| TCGACCTGTACGGGAT-1 | C198R_1M_rep2 |
| TCGTCCAAGGTAAGTT-1 | C198R_1M_rep2 |
| TCGTGGGTCACTAGCA-1 | C198R_1M_rep2 |
| TCTAACTTCCTGGCTT-1 | C198R_1M_rep2 |
| TCTCACGGTGTTCGTA-1 | C198R_1M_rep2 |
| TGCATCCCAGAGTAAT-1 | C198R_1M_rep2 |
| TTTACGTTCTATCACT-1 | C198R_1M_rep2 |
| AAGATAGCATGTACGT-1 | C198R_4M_rep1 |
| ATTCTTGCACTAGGCC-1 | C198R_4M_rep1 |
| GTGAGTTTCAAGCTGT-1 | C198R_4M_rep1 |
| TCTACATGTTGCCGCA-1 | C198R_4M_rep1 |
| TGGTGATCAAATACGA-1 | C198R_4M_rep1 |
| AAAGGATGTCGGAAAC-1 | C198R_4M_rep2 |
| AAAGTGAAGAGTGTTA-1 | C198R_4M_rep2 |
| AACAAAGCACCCTTGT-1 | C198R_4M_rep2 |
| AACACACCACTACCGG-1 | C198R_4M_rep2 |
| AACACACTCGCATAGT-1 | C198R_4M_rep2 |
| AACGGGAAGCATACTC-1 | C198R_4M_rep2 |
| AAGCCATCAAGTAGTA-1 | C198R_4M_rep2 |
| AAGCGTTCATCTCCCA-1 | C198R_4M_rep2 |
| AAGTCGTAGGGCCCTT-1 | C198R_4M_rep2 |
| AAGTTCGTCTTACCGC-1 | C198R_4M_rep2 |
| AATGCCACAAGCTGTT-1 | C198R_4M_rep2 |
| ACAAAGAAGAACTTCC-1 | C198R_4M_rep2 |
| ACACTGAGTGGGATTG-1 | C198R_4M_rep2 |
| ACATCGAGTATATGGA-1 | C198R_4M_rep2 |

| | |
|---|---|
| ACCATTTTCCTCGCAT-1 | C198R_4M_rep2 |
| ACCCTCACAGTTGTTG-1 | C198R_4M_rep2 |
| ACGATCAGTGTGGACA-1 | C198R_4M_rep2 |
| ACGGAAGCAGGACATG-1 | C198R_4M_rep2 |
| ACTATCTGTTCAATCG-1 | C198R_4M_rep2 |
| ACTTCGCCATAGGAGC-1 | C198R_4M_rep2 |
| AGAAATGGTCCTCAGG-1 | C198R_4M_rep2 |
| AGATGCTTCCTCTGCA-1 | C198R_4M_rep2 |
| AGTACCAGTTTACTTC-1 | C198R_4M_rep2 |
| AGTCAACAGCTGGTGA-1 | C198R_4M_rep2 |
| AGTCAACGTTACCGTA-1 | C198R_4M_rep2 |
| AGTCTCCTCGCCTTTG-1 | C198R_4M_rep2 |
| AGTGCCGCACTCATAG-1 | C198R_4M_rep2 |
| AGTTAGCGTGGTTTAC-1 | C198R_4M_rep2 |
| ATACCTTGTGGTCAAG-1 | C198R_4M_rep2 |
| ATACTTCGTACGCTTA-1 | C198R_4M_rep2 |
| ATAGGCTCACGTCTCT-1 | C198R_4M_rep2 |
| ATCATTCTCGTGTGGC-1 | C198R_4M_rep2 |
| ATCCGTCGTGCCGAAA-1 | C198R_4M_rep2 |
| ATCGTCCCAAATACAG-1 | C198R_4M_rep2 |
| ATCTCTATCACGGAGA-1 | C198R_4M_rep2 |
| ATGAGTCAGGTAGGCT-1 | C198R_4M_rep2 |
| ATGTCTTCAGTAGATA-1 | C198R_4M_rep2 |
| ATTCACTAGCTCGAAG-1 | C198R_4M_rep2 |
| ATTCCATCAAGTTGGG-1 | C198R_4M_rep2 |
| ATTCTTGTCGTAACTG-1 | C198R_4M_rep2 |
| ATTGGGTAGAGTCTGG-1 | C198R_4M_rep2 |
| ATTTACCGTCGTTATG-1 | C198R_4M_rep2 |
| ATTTCACGTTTCGTGA-1 | C198R_4M_rep2 |
| ATTTCTGTCTGTAAGC-1 | C198R_4M_rep2 |
| CAAAGAAAGTGGTCAG-1 | C198R_4M_rep2 |
| CAACGGCCAACGAGGT-1 | C198R_4M_rep2 |
| CAAGACTAGAGGACTC-1 | C198R_4M_rep2 |
| CAAGAGGAGTGCTCAT-1 | C198R_4M_rep2 |
| CAAGCTAAGAAGATCT-1 | C198R_4M_rep2 |
| CACAACAGTGGAACAC-1 | C198R_4M_rep2 |
| CACAGATAGTGACACG-1 | C198R_4M_rep2 |
| CACAGGCGTATGCTAC-1 | C198R_4M_rep2 |
| CACTGTCCATCGGCCA-1 | C198R_4M_rep2 |
| CAGCAATCACCTTCCA-1 | C198R_4M_rep2 |
| CATACAGCAGGTATGG-1 | C198R_4M_rep2 |
| CATGCAAAGGATATAC-1 | C198R_4M_rep2 |
| CATGGTAAGGGACTGT-1 | C198R_4M_rep2 |
| CATTCATTCGAACGCC-1 | C198R_4M_rep2 |

| | |
|---|---|
| CATTCCGGTCTTACAG-1 | C198R_4M_rep2 |
| CATTGCCGTCGCACAC-1 | C198R_4M_rep2 |
| CCACGTTCACAGACGA-1 | C198R_4M_rep2 |
| CCCATTGGTTGTTTGG-1 | C198R_4M_rep2 |
| CCCGAAGTCAAGATAG-1 | C198R_4M_rep2 |
| CCGGGTAAGGTTGTTC-1 | C198R_4M_rep2 |
| CCGTGAGGTCAAGTTC-1 | C198R_4M_rep2 |
| CCGTGAGGTTTCGATG-1 | C198R_4M_rep2 |
| CCTAACCTCACAAGGG-1 | C198R_4M_rep2 |
| CCTGTTGAGATGACCG-1 | C198R_4M_rep2 |
| CCTTGTGAGGTAAGAG-1 | C198R_4M_rep2 |
| CGAGGAAGTGAACGGT-1 | C198R_4M_rep2 |
| CGAGGCTGTCACTTAG-1 | C198R_4M_rep2 |
| CGCAGGTGTCACGCTG-1 | C198R_4M_rep2 |
| CGCCATTTCTAGCCTC-1 | C198R_4M_rep2 |
| CGGAATTAGCATAGGC-1 | C198R_4M_rep2 |
| CGGGACTCACGTATAC-1 | C198R_4M_rep2 |
| CGTCAAACAGAGAGGG-1 | C198R_4M_rep2 |
| CGTGAATGTCAGACGA-1 | C198R_4M_rep2 |
| CGTGATACATATGGCT-1 | C198R_4M_rep2 |
| CTACAGACAATGAAAC-1 | C198R_4M_rep2 |
| CTACATTCAACCACAT-1 | C198R_4M_rep2 |
| CTCACTGGTTGCAACT-1 | C198R_4M_rep2 |
| CTCAGTCTCCCATAAG-1 | C198R_4M_rep2 |
| CTCAGTCTCTGCTTTA-1 | C198R_4M_rep2 |
| CTCCGATAGACGGTCA-1 | C198R_4M_rep2 |
| CTCCGATCAAGGGTCA-1 | C198R_4M_rep2 |
| CTGAGCGAGGCCTGCT-1 | C198R_4M_rep2 |
| CTGCCATCATAATGCC-1 | C198R_4M_rep2 |
| CTGCCATTCGTGTCAA-1 | C198R_4M_rep2 |
| CTGCCTATCAGCGCGT-1 | C198R_4M_rep2 |
| CTGCGAGAGTACTGGG-1 | C198R_4M_rep2 |
| CTGGCAGCAGAACATA-1 | C198R_4M_rep2 |
| CTTACCGTCTCCCTAG-1 | C198R_4M_rep2 |
| CTTAGGATCAAGAGGC-1 | C198R_4M_rep2 |
| CTTAGGATCCGCACTT-1 | C198R_4M_rep2 |
| CTTCCGACATCCCACT-1 | C198R_4M_rep2 |
| CTTCCTTCAGCATGCC-1 | C198R_4M_rep2 |
| CTTCCTTTCCGATGTA-1 | C198R_4M_rep2 |
| GACAGCCCATAACAGA-1 | C198R_4M_rep2 |
| GACTCTCGTACACGTT-1 | C198R_4M_rep2 |
| GACTTCCGTTCTCGTC-1 | C198R_4M_rep2 |
| GAGACCCTCGGCTTGG-1 | C198R_4M_rep2 |
| GAGACTTCAGGCATGA-1 | C198R_4M_rep2 |

| | |
|---|---|
| GAGCCTGCAAGTATCC-1 | C198R_4M_rep2 |
| GAGTGTTGTCCGTTTC-1 | C198R_4M_rep2 |
| GATAGCTGTAGATGTA-1 | C198R_4M_rep2 |
| GATGGAGAGAGAGGTA-1 | C198R_4M_rep2 |
| GATGGAGGTGTCACAT-1 | C198R_4M_rep2 |
| GATTCGAAGGTCACCC-1 | C198R_4M_rep2 |
| GATTCTTGTTGTGCAT-1 | C198R_4M_rep2 |
| GCAACATGTACTAGCT-1 | C198R_4M_rep2 |
| GCCTGTTAGCGGACAT-1 | C198R_4M_rep2 |
| GCGATCGCAAGCGATG-1 | C198R_4M_rep2 |
| GCTGAATTCTGACAGT-1 | C198R_4M_rep2 |
| GGAGAACTCTTCCCGA-1 | C198R_4M_rep2 |
| GGCAGTCAGGTCACCC-1 | C198R_4M_rep2 |
| GGCTTTCAGGTCGTGA-1 | C198R_4M_rep2 |
| GGCTTTCCATAATCCG-1 | C198R_4M_rep2 |
| GGGCTACCAAATCGGG-1 | C198R_4M_rep2 |
| GGGCTACGTCGTACAT-1 | C198R_4M_rep2 |
| GGGTATTCAAGCACAG-1 | C198R_4M_rep2 |
| GGGTTTACATGTTCGA-1 | C198R_4M_rep2 |
| GGTAATCGTAGTGCGA-1 | C198R_4M_rep2 |
| GGTCTGGCAGGTCCCA-1 | C198R_4M_rep2 |
| GGTGTTACAAGCGATG-1 | C198R_4M_rep2 |
| GTATTGGGTGCCTTTC-1 | C198R_4M_rep2 |
| GTCTCACCAAGAATAC-1 | C198R_4M_rep2 |
| GTCTTTAGTATATGGA-1 | C198R_4M_rep2 |
| GTGATGTTCACTTCTA-1 | C198R_4M_rep2 |
| GTGCTTCTCCTAAGTG-1 | C198R_4M_rep2 |
| GTTGCGGAGAGAACCC-1 | C198R_4M_rep2 |
| GTTGTCCAGGCATCTT-1 | C198R_4M_rep2 |
| GTTGTGACAACGTTAC-1 | C198R_4M_rep2 |
| GTTTACTAGACTTAAG-1 | C198R_4M_rep2 |
| TAAGCCATCACAGAGG-1 | C198R_4M_rep2 |
| TAATCTCTCACCACAA-1 | C198R_4M_rep2 |
| TACAGGTAGGGCAATC-1 | C198R_4M_rep2 |
| TACTGCCAGAAGGGAT-1 | C198R_4M_rep2 |
| TACTGCCCACGTCATA-1 | C198R_4M_rep2 |
| TACTTACTCAATCCGA-1 | C198R_4M_rep2 |
| TACTTGTAGTTAGAAC-1 | C198R_4M_rep2 |
| TAGATCGAGGTTACCT-1 | C198R_4M_rep2 |
| TAGGTACAGCCGTTAT-1 | C198R_4M_rep2 |
| TATCTGTAGCGGTAAC-1 | C198R_4M_rep2 |
| TATTGCTGTCGATTTG-1 | C198R_4M_rep2 |
| TCAAGACCACTACCCT-1 | C198R_4M_rep2 |
| TCAAGCACATCCTTCG-1 | C198R_4M_rep2 |

| | |
|---|---|
| TCAAGTGGTCGTGGAA-1 | C198R_4M_rep2 |
| TCACAAGGTCCATAGT-1 | C198R_4M_rep2 |
| TCACTCGGTAGGTAGC-1 | C198R_4M_rep2 |
| TCACTCGTCACTTATC-1 | C198R_4M_rep2 |
| TCAGCAATCCGTGTCT-1 | C198R_4M_rep2 |
| TCAGTTTCAAATGAAC-1 | C198R_4M_rep2 |
| TCAGTTTCAAGAGTAT-1 | C198R_4M_rep2 |
| TCATCCGAGCATCGAG-1 | C198R_4M_rep2 |
| TCATGGAAGAGATTCA-1 | C198R_4M_rep2 |
| TCATTACTCCCTCGTA-1 | C198R_4M_rep2 |
| TCCATGCCACAGACGA-1 | C198R_4M_rep2 |
| TCCGAAATCCTTCACG-1 | C198R_4M_rep2 |
| TCCGATCCAGCACCCA-1 | C198R_4M_rep2 |
| TCCGGGAAGACACACG-1 | C198R_4M_rep2 |
| TCCTAATCACAAACGG-1 | C198R_4M_rep2 |
| TCCTCTTGTCTGTAGT-1 | C198R_4M_rep2 |
| TCGCTTGAGAGGCCAT-1 | C198R_4M_rep2 |
| TCTACATTCCTCTCTT-1 | C198R_4M_rep2 |
| TCTACCGTCGACCCAG-1 | C198R_4M_rep2 |
| TCTCCGACAGCAAGAC-1 | C198R_4M_rep2 |
| TCTTAGTCACGAGGAT-1 | C198R_4M_rep2 |
| TCTTTGAAGTGGTTAA-1 | C198R_4M_rep2 |
| TGAGCGCAGATCACCT-1 | C198R_4M_rep2 |
| TGAGGTTCAATAACGA-1 | C198R_4M_rep2 |
| TGATCAGGTAGCACGA-1 | C198R_4M_rep2 |
| TGCAGTAAGGTGCGAT-1 | C198R_4M_rep2 |
| TGCGATAAGTCACTCA-1 | C198R_4M_rep2 |
| TGCGGGTTCTTCCACG-1 | C198R_4M_rep2 |
| TGCTTGCTCCTCTCTT-1 | C198R_4M_rep2 |
| TGGGAAGCATGCAGGA-1 | C198R_4M_rep2 |
| TGGGAGAGTTCTCTCG-1 | C198R_4M_rep2 |
| TGTAACGCACCGTGAC-1 | C198R_4M_rep2 |
| TGTCAGAAGAACCCGA-1 | C198R_4M_rep2 |
| TGTCCACCATGTGCCG-1 | C198R_4M_rep2 |
| TGTGTGAAGTTCACTG-1 | C198R_4M_rep2 |
| TGTTACTTCATACGAC-1 | C198R_4M_rep2 |
| TGTTCCGAGCTGGCCT-1 | C198R_4M_rep2 |
| TGTTCCGGTGATACAA-1 | C198R_4M_rep2 |
| TGTTGGATCCGCTGTT-1 | C198R_4M_rep2 |
| TTAATCCTCAACCCGG-1 | C198R_4M_rep2 |
| TTCAATCCATCCTGTC-1 | C198R_4M_rep2 |
| TTCACGCAGACATATG-1 | C198R_4M_rep2 |
| TTCACGCTCTTTCCGG-1 | C198R_4M_rep2 |
| TTCTAGTAGAAACTCA-1 | C198R_4M_rep2 |

| | |
|---|---|
| TTGATGGCATCGTGCG-1 | C198R_4M_rep2 |
| TTGGGTATCATGGAGG-1 | C198R_4M_rep2 |
| TTGTTTGAGCTTTCCC-1 | C198R_4M_rep2 |
| TTGTTTGGTTGCTCGG-1 | C198R_4M_rep2 |
| TTTCACACAGATGCGA-1 | C198R_4M_rep2 |
| ACAGCCGGTATGACAA-1 | DKO_1M_rep1 |
| CACCAAAGTGGTCTTA-1 | DKO_1M_rep1 |
| CCAATGAGTTGCATCA-1 | DKO_1M_rep1 |
| CTACATTAGTGCGTCC-1 | DKO_1M_rep1 |
| GAAGGACTCCGACAGC-1 | DKO_1M_rep1 |
| GATGACTAGGACAGCT-1 | DKO_1M_rep1 |
| GGCTGTGGTTGTGTAC-1 | DKO_1M_rep1 |
| TCAAGACGTGACTGAG-1 | DKO_1M_rep1 |
| TCAATTCTCGGTCTAA-1 | DKO_1M_rep1 |
| TCCCATGTCATGCTAG-1 | DKO_1M_rep1 |
| AACAAAGAGTGTTCAC-1 | DKO_1M_rep2 |
| AACCATGTCTCGGGAC-1 | DKO_1M_rep2 |
| ACCTACCAGCAGGCAT-1 | DKO_1M_rep2 |
| ACGATGTCAAGCAATA-1 | DKO_1M_rep2 |
| ACGCACGGTAATCAAG-1 | DKO_1M_rep2 |
| AGGTCATTCCTTGGAA-1 | DKO_1M_rep2 |
| ATCGCCTCAGAGGCAT-1 | DKO_1M_rep2 |
| ATTACTCCATGACGTT-1 | DKO_1M_rep2 |
| ATTCAGGGTGCTGCAC-1 | DKO_1M_rep2 |
| ATTCTTGTCTTTCCAA-1 | DKO_1M_rep2 |
| CAACGGCAGAGGTTTA-1 | DKO_1M_rep2 |
| CAATACGGTCCCGTGA-1 | DKO_1M_rep2 |
| CACGTGGAGGAGTATT-1 | DKO_1M_rep2 |
| CATGCGGGTGTGACCC-1 | DKO_1M_rep2 |
| CATGGATAGTAGCTCT-1 | DKO_1M_rep2 |
| CCAATTTAGACGAGCT-1 | DKO_1M_rep2 |
| CCTCAGTCAACTGAAA-1 | DKO_1M_rep2 |
| CGTAGTACAAGTATCC-1 | DKO_1M_rep2 |
| CTGAGCGTCATTGTGG-1 | DKO_1M_rep2 |
| CTGTAGACAAGCGCAA-1 | DKO_1M_rep2 |
| CTGTAGAGTATGTCCA-1 | DKO_1M_rep2 |
| CTGTGAAGTGATATAG-1 | DKO_1M_rep2 |
| GATTCTTTCGCTATTT-1 | DKO_1M_rep2 |
| GATTTCTCACGACCTG-1 | DKO_1M_rep2 |
| GCAGTTATCGCGTCGA-1 | DKO_1M_rep2 |
| GGAAGTGAGTTTAGGA-1 | DKO_1M_rep2 |
| GGAATGGTCAGGACAG-1 | DKO_1M_rep2 |
| GGAGGTAAGTGCTCGC-1 | DKO_1M_rep2 |
| GGGTATTCACACACGC-1 | DKO_1M_rep2 |

| | |
|---|---|
| GTCACGGAGATTTGCC-1 | DKO_1M_rep2 |
| GTCAGCGTCGTCCTCA-1 | DKO_1M_rep2 |
| GTTCGCTTCATCTACT-1 | DKO_1M_rep2 |
| TACGGGCCAGTTCACA-1 | DKO_1M_rep2 |
| TAGACCACACAAGGTG-1 | DKO_1M_rep2 |
| TATTGCTAGAATCCCT-1 | DKO_1M_rep2 |
| TCAGTTTAGGGCCAAT-1 | DKO_1M_rep2 |
| TCCCATGCAGCCTATA-1 | DKO_1M_rep2 |
| TCGAACAGTTAGGAGC-1 | DKO_1M_rep2 |
| TCGACGGCAGCAGTCC-1 | DKO_1M_rep2 |
| TCGATTTAGCAACCAG-1 | DKO_1M_rep2 |
| TCTCAGCGTACATACC-1 | DKO_1M_rep2 |
| TGATGCAAGTAAAGCT-1 | DKO_1M_rep2 |
| TGCTTGCAGTTCTCTT-1 | DKO_1M_rep2 |
| TGGAGGAAGTTGTAGA-1 | DKO_1M_rep2 |
| TGTTCTAGTAAGGAGA-1 | DKO_1M_rep2 |
| TTGTGGAAGTTAGAAC-1 | DKO_1M_rep2 |
| AACCATGGTCCTACAA-1 | DKO_4M_rep1 |
| AAGCGTTGTGAATTGA-1 | DKO_4M_rep1 |
| ACAGGGATCGTTATCT-1 | DKO_4M_rep1 |
| ACTCCCACATGCCATA-1 | DKO_4M_rep1 |
| ACTGTCCAGTCCTGCG-1 | DKO_4M_rep1 |
| AGGCCACGTTGTGTTG-1 | DKO_4M_rep1 |
| ATATCCTAGGAGGTTC-1 | DKO_4M_rep1 |
| ATGGATCTCTGTCCCA-1 | DKO_4M_rep1 |
| ATTACTCAGAAACTCA-1 | DKO_4M_rep1 |
| ATTCGTTTCTGGAGAG-1 | DKO_4M_rep1 |
| CAAGAGGCACAAGCCC-1 | DKO_4M_rep1 |
| CAGTTAGAGTCTGGAG-1 | DKO_4M_rep1 |
| CATGAGTCACGAAAGC-1 | DKO_4M_rep1 |
| CCACGTTGTTCGGACC-1 | DKO_4M_rep1 |
| CCCATTGCATACTGTG-1 | DKO_4M_rep1 |
| CGGTCAGCAGGTTACT-1 | DKO_4M_rep1 |
| CTACGGGGTCCTTAAG-1 | DKO_4M_rep1 |
| CTCACTGTCTGGTCAA-1 | DKO_4M_rep1 |
| GAGGGATCAGAGGCTA-1 | DKO_4M_rep1 |
| GATCACAGTCTTCATT-1 | DKO_4M_rep1 |
| GCACGGTCATACTTTC-1 | DKO_4M_rep1 |
| GGAACCCTCTTAGCTT-1 | DKO_4M_rep1 |
| GGAATGGGTGCTGTCG-1 | DKO_4M_rep1 |
| GTCTTTATCAAGGACG-1 | DKO_4M_rep1 |
| GTTTGGAAGCCTATCA-1 | DKO_4M_rep1 |
| TACCCGTTCGAGCCTG-1 | DKO_4M_rep1 |
| TACGGTATCTCAGGCG-1 | DKO_4M_rep1 |

| | |
|---|---|
| TCAAGACAGACGTCGA-1 | DKO_4M_rep1 |
| TCATCCGGTGTTGACT-1 | DKO_4M_rep1 |
| TCCCAGTCAAGGCGTA-1 | DKO_4M_rep1 |
| TCTTCCTCATTCGGGC-1 | DKO_4M_rep1 |
| TGAACGTGTGAAGCGT-1 | DKO_4M_rep1 |
| TGAGCATAGAGCCTGA-1 | DKO_4M_rep1 |
| TGCATGATCCGCAAAT-1 | DKO_4M_rep1 |
| TGTTCTAGTCAAGTTC-1 | DKO_4M_rep1 |
| TGTTTGTTCCATTTGT-1 | DKO_4M_rep1 |
| TTAGGCAAGATAGGGA-1 | DKO_4M_rep1 |
| TTGCTGCGTGGCTCTG-1 | DKO_4M_rep1 |
| TTGGATGCAGTGTATC-1 | DKO_4M_rep1 |
| TTTCCTCCAGAACGCA-1 | DKO_4M_rep1 |
| TTTGGTTTCGACATAC-1 | DKO_4M_rep1 |
| ACGTTCCTCTATCCAT-1 | DKO_4M_rep2 |
| AGTCACATCTCTCTTC-1 | DKO_4M_rep2 |
| GATCAGTCACCAGACC-1 | DKO_4M_rep2 |
| TAGTGCAAGAGGTTTA-1 | DKO_4M_rep2 |
| TGACGCGGTCTTGCGG-1 | DKO_4M_rep2 |
| AACACACGTTCTAACG-1 | WT_1M_rep1 |
| AACGTCATCCTATTTG-1 | WT_1M_rep1 |
| AATGCCATCCCAGGCA-1 | WT_1M_rep1 |
| ATGCGATAGACGTCCC-1 | WT_1M_rep1 |
| CATAAGCAGGAAGAAC-1 | WT_1M_rep1 |
| CATGCTCAGCCATTTG-1 | WT_1M_rep1 |
| CCGGGTATCTCCGAGG-1 | WT_1M_rep1 |
| CGATGGCTCCTTGACC-1 | WT_1M_rep1 |
| CGCCATTCAGAAATCA-1 | WT_1M_rep1 |
| CGGAGAACATAGACTC-1 | WT_1M_rep1 |
| CGGGACTTCGTGCGAC-1 | WT_1M_rep1 |
| CGTGCTTTCTAGAACC-1 | WT_1M_rep1 |
| CTCAAGACACGCTTAA-1 | WT_1M_rep1 |
| CTCCGATAGATACTGA-1 | WT_1M_rep1 |
| GAAGGGTGTCTGTCAA-1 | WT_1M_rep1 |
| GAGCCTGAGTTGGCGA-1 | WT_1M_rep1 |
| GAGCCTGTCGACCATA-1 | WT_1M_rep1 |
| GAGTCATAGCGTGCCT-1 | WT_1M_rep1 |
| GATCGTAAGAAGCGAA-1 | WT_1M_rep1 |
| GGGATCCGTACCTGTA-1 | WT_1M_rep1 |
| GGGTTATCAAACGGCA-1 | WT_1M_rep1 |
| GGTAATCAGGATACCG-1 | WT_1M_rep1 |
| GTCAAGTCAATAGGGC-1 | WT_1M_rep1 |
| GTTGCGGTCCGGTTCT-1 | WT_1M_rep1 |
| TACTTACCATCGCTCT-1 | WT_1M_rep1 |

| | |
|---|---|
| TCACACCGTTGCAACT-1 | WT_1M_rep1 |
| TCACTATAGCCTCGTG-1 | WT_1M_rep1 |
| TCTCAGCAGGGCAAGG-1 | WT_1M_rep1 |
| TCTCTGGTCCCATTTA-1 | WT_1M_rep1 |
| TGGCGTGCAGGTGTGA-1 | WT_1M_rep1 |
| TGTTGAGAGAGGCTGT-1 | WT_1M_rep1 |
| TTAGTCTCAGATTTCG-1 | WT_1M_rep1 |
| TTCTCTCTCCGCACTT-1 | WT_1M_rep1 |
| TTGCCTGGTTGATCGT-1 | WT_1M_rep1 |
| TTTAGTCAGGTTAGTA-1 | WT_1M_rep1 |
| TTTGGTTCATATCTCT-1 | WT_1M_rep1 |
| AACCTTTAGCTCATAC-1 | WT_1M_rep2 |
| ACGATCACACAATTCG-1 | WT_1M_rep2 |
| AGGGTCCAGTTGGGAC-1 | WT_1M_rep2 |
| ATTTCACCACTTCAGA-1 | WT_1M_rep2 |
| CTCCCTCAGAGCCTGA-1 | WT_1M_rep2 |
| TCACTCGAGAATTTGG-1 | WT_1M_rep2 |
| TTCTTGATCCAGGACC-1 | WT_1M_rep2 |
| TTTACTGGTCCGAAGA-1 | WT_1M_rep2 |
| AACAACCCAGGACGAT-1 | WT_4M_rep1 |
| AACAAGAGTACAATAG-1 | WT_4M_rep1 |
| AACAGGGTCAAGATAG-1 | WT_4M_rep1 |
| AACCACACATCTTTCA-1 | WT_4M_rep1 |
| AACCACAGTCTCAAGT-1 | WT_4M_rep1 |
| AACCTGAGTGTGTCGC-1 | WT_4M_rep1 |
| AACCTTTTCGGTATGT-1 | WT_4M_rep1 |
| AAGCCATGTGACAGCA-1 | WT_4M_rep1 |
| AAGCCATTCTTACCGC-1 | WT_4M_rep1 |
| AAGCGAGCATATGCGT-1 | WT_4M_rep1 |
| AAGTACCCAGACGATG-1 | WT_4M_rep1 |
| AAGTGAAAGCTAATGA-1 | WT_4M_rep1 |
| AATCACGAGTCAGGGT-1 | WT_4M_rep1 |
| AATGGAACAGTATTCG-1 | WT_4M_rep1 |
| AATTCCTGTTGAATCC-1 | WT_4M_rep1 |
| AATTCCTTCAGCTAGT-1 | WT_4M_rep1 |
| ACAAGCTCATTCACAG-1 | WT_4M_rep1 |
| ACACTGAGTGCACGCT-1 | WT_4M_rep1 |
| ACAGAAACATTGGGAG-1 | WT_4M_rep1 |
| ACATGCAGTTAAACCC-1 | WT_4M_rep1 |
| ACATTTCCAAGTCGTT-1 | WT_4M_rep1 |
| ACCAACATCTTCCAGC-1 | WT_4M_rep1 |
| ACCCAAAGTGGATGAC-1 | WT_4M_rep1 |
| ACCCTCATCTTTCCGG-1 | WT_4M_rep1 |
| ACCTGAAGTATCTTCT-1 | WT_4M_rep1 |

| | |
|---|---|
| ACCTGTCTCGAATGCT-1 | WT_4M_rep1 |
| ACGATCACACAAATAG-1 | WT_4M_rep1 |
| ACGATGTCAATCTCTT-1 | WT_4M_rep1 |
| ACGGAAGAGAAGTGTT-1 | WT_4M_rep1 |
| ACGGAAGGTCGTATGT-1 | WT_4M_rep1 |
| ACGGGTCAGAGTATAC-1 | WT_4M_rep1 |
| ACGTACAAGAAGCTCG-1 | WT_4M_rep1 |
| ACGTACAAGTCTAGCT-1 | WT_4M_rep1 |
| ACGTAGTCAGCAGTCC-1 | WT_4M_rep1 |
| ACGTCCTTCTCTGGTC-1 | WT_4M_rep1 |
| ACGTTCCCAGCAATTC-1 | WT_4M_rep1 |
| ACTACGACAAATCGTC-1 | WT_4M_rep1 |
| ACTATGGAGTAGGCCA-1 | WT_4M_rep1 |
| ACTATTCTCTGTGCTC-1 | WT_4M_rep1 |
| ACTGCAAAGAATAGTC-1 | WT_4M_rep1 |
| ACTGCAATCGAGTCTA-1 | WT_4M_rep1 |
| AGACACTGTGGACTAG-1 | WT_4M_rep1 |
| AGACAGGCACTGCGTG-1 | WT_4M_rep1 |
| AGACAGGGTACGCGTC-1 | WT_4M_rep1 |
| AGATAGATCGACGACC-1 | WT_4M_rep1 |
| AGCGTATTCGCTTGAA-1 | WT_4M_rep1 |
| AGGAATATCCATAGAC-1 | WT_4M_rep1 |
| AGGACGACATCACGGC-1 | WT_4M_rep1 |
| AGGATAATCAACCTCC-1 | WT_4M_rep1 |
| AGGATAATCCATTTAC-1 | WT_4M_rep1 |
| AGGATCTAGGTTATAG-1 | WT_4M_rep1 |
| AGGCTGCCAGACATCT-1 | WT_4M_rep1 |
| AGGGTTTTCACTCACC-1 | WT_4M_rep1 |
| AGGTCATGTAAGGAGA-1 | WT_4M_rep1 |
| AGGTGTTTCTCACGAA-1 | WT_4M_rep1 |
| AGGTTGTGTGTACATC-1 | WT_4M_rep1 |
| AGTAACCTCATAGAGA-1 | WT_4M_rep1 |
| AGTAACCTCGTGTTCC-1 | WT_4M_rep1 |
| AGTAGCTGTTAATGAG-1 | WT_4M_rep1 |
| AGTCACACAGGTCAAG-1 | WT_4M_rep1 |
| AGTGTTGCATCCGGCA-1 | WT_4M_rep1 |
| ATACCGACACAATGTC-1 | WT_4M_rep1 |
| ATACCGACACGATTCA-1 | WT_4M_rep1 |
| ATACTTCAGGATCACG-1 | WT_4M_rep1 |
| ATAGGCTGTGAAGCTG-1 | WT_4M_rep1 |
| ATCACAGTCGCTCCTA-1 | WT_4M_rep1 |
| ATCACGAAGTTCTCTT-1 | WT_4M_rep1 |
| ATCAGGTAGATGGTAT-1 | WT_4M_rep1 |
| ATCCCTGTCCGTTGAA-1 | WT_4M_rep1 |

| | |
|---|---|
| ATCGCCTTCAACACCA-1 | WT_4M_rep1 |
| ATCGTCCTCAACTTTC-1 | WT_4M_rep1 |
| ATGACCAAGGGTTTCT-1 | WT_4M_rep1 |
| ATGGAGGCAGCGTGCT-1 | WT_4M_rep1 |
| ATGGAGGGTGCCAAGA-1 | WT_4M_rep1 |
| ATGGAGGTCATGCCGG-1 | WT_4M_rep1 |
| ATGGGTTAGTATTAGG-1 | WT_4M_rep1 |
| ATGGGTTCACGTGAGA-1 | WT_4M_rep1 |
| ATTACCTGTGTGTACT-1 | WT_4M_rep1 |
| ATTCAGGCAATAGAGT-1 | WT_4M_rep1 |
| ATTTCTGAGTACAACA-1 | WT_4M_rep1 |
| CAAGAGGAGCGTTGTT-1 | WT_4M_rep1 |
| CAAGCTATCGCGTAGC-1 | WT_4M_rep1 |
| CAAGGGAGTCACAATC-1 | WT_4M_rep1 |
| CAAGGGATCACAAGAA-1 | WT_4M_rep1 |
| CAATACGCACTTTATC-1 | WT_4M_rep1 |
| CAATGACAGGCAGCTA-1 | WT_4M_rep1 |
| CAATTTCCAAGACCTT-1 | WT_4M_rep1 |
| CACACAAAGGCCACTC-1 | WT_4M_rep1 |
| CACAGATCATTCCTAT-1 | WT_4M_rep1 |
| CACATGAGTCCTGGGT-1 | WT_4M_rep1 |
| CACCAAACAGAGTTGG-1 | WT_4M_rep1 |
| CACCGTTGTACCTAGT-1 | WT_4M_rep1 |
| CACGAATAGCTAGATA-1 | WT_4M_rep1 |
| CACGAATTCATCACAG-1 | WT_4M_rep1 |
| CAGGTATCAATAGTGA-1 | WT_4M_rep1 |
| CAGTGCGAGCACTTTG-1 | WT_4M_rep1 |
| CAGTTAGTCCATATGG-1 | WT_4M_rep1 |
| CATAAGCCAAGGTCGA-1 | WT_4M_rep1 |
| CATACCCAGAGGCGTT-1 | WT_4M_rep1 |
| CATCCCATCGCCAACG-1 | WT_4M_rep1 |
| CATTCATTCCTTCTTC-1 | WT_4M_rep1 |
| CATTCCGAGAGTCCGA-1 | WT_4M_rep1 |
| CATTCTACATGCAGGA-1 | WT_4M_rep1 |
| CATTGCCGTAGCACAG-1 | WT_4M_rep1 |
| CCAATGAAGTACAGAT-1 | WT_4M_rep1 |
| CCACAAATCTAGCCAA-1 | WT_4M_rep1 |
| CCACGTTTCCCTATTA-1 | WT_4M_rep1 |
| CCCAACTGTCCTGTTC-1 | WT_4M_rep1 |
| CCCATTGAGTCATGGG-1 | WT_4M_rep1 |
| CCCATTGAGTGGTTAA-1 | WT_4M_rep1 |
| CCCTCTCAGTCGGCCT-1 | WT_4M_rep1 |
| CCGCAAGAGTCTAGCT-1 | WT_4M_rep1 |
| CCGTGAGCACATTCTT-1 | WT_4M_rep1 |

| | |
|---|---|
| CCGTTCAAGCTGAAAT-1 | WT_4M_rep1 |
| CCGTTCAGTTCAGCGC-1 | WT_4M_rep1 |
| CCTAAGACACGGATCC-1 | WT_4M_rep1 |
| CCTATCGGTATCGTGT-1 | WT_4M_rep1 |
| CCTCAGTTCGCCGAAC-1 | WT_4M_rep1 |
| CCTCCAAGTTCCACAA-1 | WT_4M_rep1 |
| CCTCCTCTCGCAAGAG-1 | WT_4M_rep1 |
| CCTCTCCAGCACGGAT-1 | WT_4M_rep1 |
| CCTTCAGCATATCGGT-1 | WT_4M_rep1 |
| CGAAGTTGTATCCTCC-1 | WT_4M_rep1 |
| CGACAGCTCCGCACGA-1 | WT_4M_rep1 |
| CGAGTTAGTACTAAGA-1 | WT_4M_rep1 |
| CGATCGGTCCCTCTCC-1 | WT_4M_rep1 |
| CGCCATTCACGTAGTT-1 | WT_4M_rep1 |
| CGGGACTGTGTAGGAC-1 | WT_4M_rep1 |
| CGGGCATAGAGCATTA-1 | WT_4M_rep1 |
| CGTAAGTGTCCTCCTA-1 | WT_4M_rep1 |
| CGTAATGAGCCTGTGC-1 | WT_4M_rep1 |
| CGTGATAAGCTTCTAG-1 | WT_4M_rep1 |
| CTAACTTAGGGACCAT-1 | WT_4M_rep1 |
| CTAAGTGTCGTGGGAA-1 | WT_4M_rep1 |
| CTAGGTACAAAGACTA-1 | WT_4M_rep1 |
| CTATCCGTCGTGGTAT-1 | WT_4M_rep1 |
| CTCAACCTCTTGATTC-1 | WT_4M_rep1 |
| CTCAATTGTCAATGGG-1 | WT_4M_rep1 |
| CTCAGGGGTCAGACGA-1 | WT_4M_rep1 |
| CTCCCTCAGTTTCAGC-1 | WT_4M_rep1 |
| CTCCGATCATTGTAGC-1 | WT_4M_rep1 |
| CTCCTTTCAACGGCCT-1 | WT_4M_rep1 |
| CTCTGGTCATACAGCT-1 | WT_4M_rep1 |
| CTGATCCAGCTATCCA-1 | WT_4M_rep1 |
| CTGCATCAGCCATATC-1 | WT_4M_rep1 |
| CTGCATCCAGCGATTT-1 | WT_4M_rep1 |
| CTGCCTAAGTCGAAAT-1 | WT_4M_rep1 |
| CTGCTCAGTGTTAACC-1 | WT_4M_rep1 |
| CTGTAGAAGATGCGAC-1 | WT_4M_rep1 |
| CTGTGAAGTATTCTCT-1 | WT_4M_rep1 |
| CTGTGAAGTTCAAGGG-1 | WT_4M_rep1 |
| CTGTGGGCAACCACGC-1 | WT_4M_rep1 |
| CTTCTAACATGAGTAA-1 | WT_4M_rep1 |
| CTTCTCTTCACTGTTT-1 | WT_4M_rep1 |
| CTTTCGGTCATCACAG-1 | WT_4M_rep1 |
| GAACACTCAACTGAAA-1 | WT_4M_rep1 |
| GAACTGTAGGTGGTTG-1 | WT_4M_rep1 |

| | |
|---|---|
| GAAGTAAGTGTCCGGT-1 | WT_4M_rep1 |
| GAATAGACAATATCCG-1 | WT_4M_rep1 |
| GACAGCCGTTAGAAAC-1 | WT_4M_rep1 |
| GACGTTAAGCATATGA-1 | WT_4M_rep1 |
| GACGTTAGTCAAGGCA-1 | WT_4M_rep1 |
| GACTATGAGGTTTACC-1 | WT_4M_rep1 |
| GACTATGTCCAAACCA-1 | WT_4M_rep1 |
| GACTCTCCACTTGGGC-1 | WT_4M_rep1 |
| GAGAGGTCATGGCTAT-1 | WT_4M_rep1 |
| GAGCCTGCACCTTCGT-1 | WT_4M_rep1 |
| GAGGCAAGTGGCTTGC-1 | WT_4M_rep1 |
| GAGTTTGTCCTAGCTC-1 | WT_4M_rep1 |
| GATAGCTCACTCATAG-1 | WT_4M_rep1 |
| GATCAGTCAGAGCTAG-1 | WT_4M_rep1 |
| GATGACTGTCGGAAAC-1 | WT_4M_rep1 |
| GATGCTAGTGGTCTCG-1 | WT_4M_rep1 |
| GCAGTTAGTAGAGATT-1 | WT_4M_rep1 |
| GCCCAGAGTGTAGCAG-1 | WT_4M_rep1 |
| GCTCAAAAGGGAACAA-1 | WT_4M_rep1 |
| GCTTCACCACAGAGAC-1 | WT_4M_rep1 |
| GCTTTCGCATATGGCT-1 | WT_4M_rep1 |
| GGAACCCAGCCTCACG-1 | WT_4M_rep1 |
| GGAAGTGTCCGGTAAT-1 | WT_4M_rep1 |
| GGAGATGTCAGCTTGA-1 | WT_4M_rep1 |
| GGAGATGTCCCAGGCA-1 | WT_4M_rep1 |
| GGCTTTCGTATTGAGA-1 | WT_4M_rep1 |
| GGGACCTGTTCGTTCC-1 | WT_4M_rep1 |
| GGGATGAAGCATACTC-1 | WT_4M_rep1 |
| GGGATGATCGTAACTG-1 | WT_4M_rep1 |
| GGGTGTCGTGGTACAG-1 | WT_4M_rep1 |
| GGTAACTTCAAAGAAC-1 | WT_4M_rep1 |
| GGTAATCAGGCCACTC-1 | WT_4M_rep1 |
| GGTGATTGTGAGCCAA-1 | WT_4M_rep1 |
| GGTGTTAAGTTAACAG-1 | WT_4M_rep1 |
| GGTTAACAGAAGTGTT-1 | WT_4M_rep1 |
| GTAAGTCGTGCAATAA-1 | WT_4M_rep1 |
| GTAGCTAGTCTACACA-1 | WT_4M_rep1 |
| GTAGTACCAAATTAGG-1 | WT_4M_rep1 |
| GTCAAACAGCTGACAG-1 | WT_4M_rep1 |
| GTCATTTTCTTTGATC-1 | WT_4M_rep1 |
| GTCGTTCCACTGATTG-1 | WT_4M_rep1 |
| GTCTCACAGCTTGTTG-1 | WT_4M_rep1 |
| GTCTTTAAGACGGTTG-1 | WT_4M_rep1 |
| GTGGGAAGTCCTTTGC-1 | WT_4M_rep1 |

| | |
|---|---|
| GTGTAACTCCAGCACG-1 | WT_4M_rep1 |
| GTGTGATAGCGTTAGG-1 | WT_4M_rep1 |
| GTGTTCCCAAGCCATT-1 | WT_4M_rep1 |
| GTGTTCCTCTCTGGTC-1 | WT_4M_rep1 |
| GTTACAGAGGAGCTGT-1 | WT_4M_rep1 |
| GTTACCCAGGTAAAGG-1 | WT_4M_rep1 |
| GTTACCCTCTGTCAGA-1 | WT_4M_rep1 |
| GTTCCGTGTACAGGTG-1 | WT_4M_rep1 |
| GTTGAACTCTAGCAAC-1 | WT_4M_rep1 |
| GTTGCTCGTAGCTTAC-1 | WT_4M_rep1 |
| GTTGTAGAGGTGCAGT-1 | WT_4M_rep1 |
| GTTTACTAGTTCTCTT-1 | WT_4M_rep1 |
| GTTTACTGTATCGTTG-1 | WT_4M_rep1 |
| TAACCAGCAGCTCCTT-1 | WT_4M_rep1 |
| TACAACGGTTCTTAGG-1 | WT_4M_rep1 |
| TACCCACCATGCCGCA-1 | WT_4M_rep1 |
| TACCCGTTCCTTCTGG-1 | WT_4M_rep1 |
| TACCTCGTCACTGGTA-1 | WT_4M_rep1 |
| TACCTGCTCTGTTGGA-1 | WT_4M_rep1 |
| TACGGGCAGTCTGCGC-1 | WT_4M_rep1 |
| TACGGGCCAGCCGTCA-1 | WT_4M_rep1 |
| TACGGTACAATAACCC-1 | WT_4M_rep1 |
| TACTGCCCAAAGGGCT-1 | WT_4M_rep1 |
| TAGACTGGTTATGGTC-1 | WT_4M_rep1 |
| TAGGAGGGTCGCTCGA-1 | WT_4M_rep1 |
| TAGGTACTCACTGATG-1 | WT_4M_rep1 |
| TATACCTTCAAGCCCG-1 | WT_4M_rep1 |
| TATCTTGGTGCCGTAC-1 | WT_4M_rep1 |
| TATTGGGGTCTACGAT-1 | WT_4M_rep1 |
| TCAAGACAGCCTCTTC-1 | WT_4M_rep1 |
| TCAAGCAGTTCGGCTG-1 | WT_4M_rep1 |
| TCACACCAGAGCTGAC-1 | WT_4M_rep1 |
| TCACTCGTCCACGGGT-1 | WT_4M_rep1 |
| TCAGTGATCCGAGGCT-1 | WT_4M_rep1 |
| TCATGTTAGCACTCAT-1 | WT_4M_rep1 |
| TCATTCATCATTTGTC-1 | WT_4M_rep1 |
| TCCATCGAGTGACCTT-1 | WT_4M_rep1 |
| TCCGAAAGTAACATGA-1 | WT_4M_rep1 |
| TCCGATCCATCTGCGG-1 | WT_4M_rep1 |
| TCCTCTTAGTATAGAC-1 | WT_4M_rep1 |
| TCGAACATCCGTAGGC-1 | WT_4M_rep1 |
| TCGATTTGTTGGAGGT-1 | WT_4M_rep1 |
| TCGCACTAGCCGATAG-1 | WT_4M_rep1 |
| TCGCAGGCAGATTAAG-1 | WT_4M_rep1 |

| | |
|---|---|
| TCGGGCAGTCAACCTA-1 | WT_4M_rep1 |
| TCTCACGAGCGTTAGG-1 | WT_4M_rep1 |
| TCTCACGCAGCAGTAG-1 | WT_4M_rep1 |
| TCTGCCACATTAGGCT-1 | WT_4M_rep1 |
| TCTGCCATCCCTATTA-1 | WT_4M_rep1 |
| TGAACGTGTTGTGCAT-1 | WT_4M_rep1 |
| TGAATGCTCAGTCAGT-1 | WT_4M_rep1 |
| TGACCCTGTATAATGG-1 | WT_4M_rep1 |
| TGACTCCAGTTGCCCG-1 | WT_4M_rep1 |
| TGAGCATCACGACGAA-1 | WT_4M_rep1 |
| TGAGCATGTGCATTAC-1 | WT_4M_rep1 |
| TGAGGTTTCAGCCTCT-1 | WT_4M_rep1 |
| TGAGTCACAAATACGA-1 | WT_4M_rep1 |
| TGAGTCATCATAGAGA-1 | WT_4M_rep1 |
| TGATCAGCATGACTTG-1 | WT_4M_rep1 |
| TGATTCTCACCATTCC-1 | WT_4M_rep1 |
| TGCACGGAGTTGGCTT-1 | WT_4M_rep1 |
| TGCATCCTCACGGGAA-1 | WT_4M_rep1 |
| TGCGGGTAGTGGCGAT-1 | WT_4M_rep1 |
| TGCTGAAAGGTGCCAA-1 | WT_4M_rep1 |
| TGCTTCGTCAACTCTT-1 | WT_4M_rep1 |
| TGGCGTGTCGACCATA-1 | WT_4M_rep1 |
| TGGGAAGTCCGATAGT-1 | WT_4M_rep1 |
| TGGGAGATCACTGATG-1 | WT_4M_rep1 |
| TGGTAGTAGCTGACTT-1 | WT_4M_rep1 |
| TGGTTAGGTTACGGAG-1 | WT_4M_rep1 |
| TGTAAGCCAACACAGG-1 | WT_4M_rep1 |
| TGTACAGCATGCGGTC-1 | WT_4M_rep1 |
| TGTGATGTCCGTGGGT-1 | WT_4M_rep1 |
| TGTGTGACACAAATCC-1 | WT_4M_rep1 |
| TGTTTGTAGACTCAAA-1 | WT_4M_rep1 |
| TTACAGGAGCGGGTTA-1 | WT_4M_rep1 |
| TTACCATGTTGTAGCT-1 | WT_4M_rep1 |
| TTAGGCAGTTGCACGC-1 | WT_4M_rep1 |
| TTCACGCGTATCTTCT-1 | WT_4M_rep1 |
| TTCATTGTCTATGTGG-1 | WT_4M_rep1 |
| TTCCACGAGATAACGT-1 | WT_4M_rep1 |
| TTCCGGTAGGGCAAGG-1 | WT_4M_rep1 |
| TTCCGGTGTGACCTGC-1 | WT_4M_rep1 |
| TTCCTCTCAAGTTCCA-1 | WT_4M_rep1 |
| TTCCTTCCAGGTCAAG-1 | WT_4M_rep1 |
| TTCTAACCACAAATGA-1 | WT_4M_rep1 |
| TTCTCTCCAGTCGTTA-1 | WT_4M_rep1 |
| TTGCGTCGTCACCCTT-1 | WT_4M_rep1 |

| | |
|---|---|
| TTGTGTTGTTTGTTCT-1 | WT_4M_rep1 |
| TTTATGCAGGAGGTTC-1 | WT_4M_rep1 |
| TTTCACAAGGCCCGTT-1 | WT_4M_rep1 |
| TTTCCTCCAAGTGGGT-1 | WT_4M_rep1 |
| TTTGACTGTAGTCTGT-1 | WT_4M_rep1 |
| TTTGTTGGTCGCACAC-1 | WT_4M_rep1 |
| AAACGAATCGGTGAAG-1 | WT_4M_rep2 |
| AAATGGACAGTGGCTC-1 | WT_4M_rep2 |
| AACAAGATCCTACAAG-1 | WT_4M_rep2 |
| AACCTTTCACAACGTT-1 | WT_4M_rep2 |
| AACTTCTTCTGTCGCT-1 | WT_4M_rep2 |
| AAGACAACATTCAGCA-1 | WT_4M_rep2 |
| AAGACTCGTAAGCAAT-1 | WT_4M_rep2 |
| AAGACTCGTAATACCC-1 | WT_4M_rep2 |
| AAGCATCAGATGGTCG-1 | WT_4M_rep2 |
| AAGCCATTCGTGTGAT-1 | WT_4M_rep2 |
| AAGTCGTCATAGCACT-1 | WT_4M_rep2 |
| AAGTGAACACTGTGAT-1 | WT_4M_rep2 |
| AAGTGAATCAGGACAG-1 | WT_4M_rep2 |
| AATAGAGCATAGGCGA-1 | WT_4M_rep2 |
| AATCGACTCAGCGTCG-1 | WT_4M_rep2 |
| AATCGTGAGCACCGTC-1 | WT_4M_rep2 |
| AATGACCAGCTAGAGC-1 | WT_4M_rep2 |
| ACAGCCGAGTAGGAAG-1 | WT_4M_rep2 |
| ACATGCACATGACGGA-1 | WT_4M_rep2 |
| ACATGCATCTTTGGAG-1 | WT_4M_rep2 |
| ACATTTCAGCCTCCAG-1 | WT_4M_rep2 |
| ACCAACACACTAGGTT-1 | WT_4M_rep2 |
| ACCAACATCCCGAACG-1 | WT_4M_rep2 |
| ACCACAACAACGTAAA-1 | WT_4M_rep2 |
| ACGATGTCACCAAATC-1 | WT_4M_rep2 |
| ACGGTTAGTTAAGGAT-1 | WT_4M_rep2 |
| ACGTCCTCAAGTGTCT-1 | WT_4M_rep2 |
| ACTACGAAGGACTGGT-1 | WT_4M_rep2 |
| ACTATCTTCATGGTAC-1 | WT_4M_rep2 |
| ACTCCCACAGACGCTC-1 | WT_4M_rep2 |
| ACTTTCATCCACACCT-1 | WT_4M_rep2 |
| ACTTTGTAGTAACCGG-1 | WT_4M_rep2 |
| AGAACAACAATGACCT-1 | WT_4M_rep2 |
| AGAACAATCGTGAGAG-1 | WT_4M_rep2 |
| AGAACCTCACAGTATC-1 | WT_4M_rep2 |
| AGAAGTATCGGTTGTA-1 | WT_4M_rep2 |
| AGAAGTATCTGAGTCA-1 | WT_4M_rep2 |
| AGACAAACAAGCCATT-1 | WT_4M_rep2 |

| | |
|---|---|
| AGAGAATAGGTGAGAA-1 | WT_4M_rep2 |
| AGAGAGCGTCCCTCAT-1 | WT_4M_rep2 |
| AGAGCAGAGGTGCGAT-1 | WT_4M_rep2 |
| AGCGATTCACTAGGTT-1 | WT_4M_rep2 |
| AGCGCCATCACCTGGG-1 | WT_4M_rep2 |
| AGCGCTGAGCGGATCA-1 | WT_4M_rep2 |
| AGCGCTGGTACCTAAC-1 | WT_4M_rep2 |
| AGCTACACATTGAAGA-1 | WT_4M_rep2 |
| AGGAATACATCCTAAG-1 | WT_4M_rep2 |
| AGGATCTTCGCCCAGA-1 | WT_4M_rep2 |
| AGGCTGCCACATATGC-1 | WT_4M_rep2 |
| AGGCTGCTCTCGCAGG-1 | WT_4M_rep2 |
| AGGGAGTTCCTAGAGT-1 | WT_4M_rep2 |
| AGGGTCCTCACAACCA-1 | WT_4M_rep2 |
| AGGGTGAAGTCTCCTC-1 | WT_4M_rep2 |
| AGGTTACAGGCGTCCT-1 | WT_4M_rep2 |
| AGGTTGTGTATGCGGA-1 | WT_4M_rep2 |
| AGGTTGTTCCATCCGT-1 | WT_4M_rep2 |
| AGTGCCGAGTGGTTAA-1 | WT_4M_rep2 |
| AGTTCCCCATGACTCA-1 | WT_4M_rep2 |
| AGTTCGAAGGCGAACT-1 | WT_4M_rep2 |
| AGTTCGAGTTTCCAAG-1 | WT_4M_rep2 |
| ATACCTTCAGAGAGGG-1 | WT_4M_rep2 |
| ATACTTCTCTGTGCTC-1 | WT_4M_rep2 |
| ATAGAGAAGGTTATAG-1 | WT_4M_rep2 |
| ATAGGCTGTCACTCGG-1 | WT_4M_rep2 |
| ATCACGAAGTGCAACG-1 | WT_4M_rep2 |
| ATCATTCTCGAGAAAT-1 | WT_4M_rep2 |
| ATCATTCTCGGATAAA-1 | WT_4M_rep2 |
| ATCCACCAGAGAGCAA-1 | WT_4M_rep2 |
| ATCGTAGTCGACGATT-1 | WT_4M_rep2 |
| ATCGTGACACAGCTGC-1 | WT_4M_rep2 |
| ATCGTGAGTGATACTC-1 | WT_4M_rep2 |
| ATCTCTATCGTGACTA-1 | WT_4M_rep2 |
| ATGAGTCAGTATGGAT-1 | WT_4M_rep2 |
| ATGAGTCCAGCGACAA-1 | WT_4M_rep2 |
| ATGAGTCGTTCGGTAT-1 | WT_4M_rep2 |
| ATTACCTTCACCTTGC-1 | WT_4M_rep2 |
| ATTACCTTCGGCTGAC-1 | WT_4M_rep2 |
| ATTATCCGTTCATCTT-1 | WT_4M_rep2 |
| ATTCATCCATCATCTT-1 | WT_4M_rep2 |
| ATTCCTACAGCATCTA-1 | WT_4M_rep2 |
| ATTCTACAGAAGGTAG-1 | WT_4M_rep2 |
| ATTGTTCGTCTTCCGT-1 | WT_4M_rep2 |

| | |
|---|---|
| CAACCTCAGTATTCCG-1 | WT_4M_rep2 |
| CAACCTCTCGACCTAA-1 | WT_4M_rep2 |
| CAACGGCCAGCGTGCT-1 | WT_4M_rep2 |
| CAACGGCGTGGCGTAA-1 | WT_4M_rep2 |
| CAATGACAGAAATTCG-1 | WT_4M_rep2 |
| CAATTTCAGGACGCTA-1 | WT_4M_rep2 |
| CACATGAAGACATCAA-1 | WT_4M_rep2 |
| CACGGGTCACTTGTGA-1 | WT_4M_rep2 |
| CACTGAACATGTGTCA-1 | WT_4M_rep2 |
| CACTGAAGTACATACC-1 | WT_4M_rep2 |
| CACTTCGTCACCTACC-1 | WT_4M_rep2 |
| CAGGCCACAAACTAAG-1 | WT_4M_rep2 |
| CAGGCCATCGAAACAA-1 | WT_4M_rep2 |
| CATACTTTCATGCATG-1 | WT_4M_rep2 |
| CATCCCAAGGCATCGA-1 | WT_4M_rep2 |
| CATGAGTAGAGTATAC-1 | WT_4M_rep2 |
| CATTCATGTCTAATCG-1 | WT_4M_rep2 |
| CATTGAGCACAGCTTA-1 | WT_4M_rep2 |
| CCAAGCGAGGAGAGGC-1 | WT_4M_rep2 |
| CCAAGCGTCTCCGTGT-1 | WT_4M_rep2 |
| CCAATGAGTAGGATAT-1 | WT_4M_rep2 |
| CCAATGATCCAAGAGG-1 | WT_4M_rep2 |
| CCAATGATCTTCTTCC-1 | WT_4M_rep2 |
| CCACGTTAGGTGTGAC-1 | WT_4M_rep2 |
| CCCTAACGTAGTGGCA-1 | WT_4M_rep2 |
| CCCTGATTCTTCTAAC-1 | WT_4M_rep2 |
| CCCTTAGAGAATTGTG-1 | WT_4M_rep2 |
| CCGAACGCACATACGT-1 | WT_4M_rep2 |
| CCGCAAGGTGGTTTGT-1 | WT_4M_rep2 |
| CCGGACATCAACGTGT-1 | WT_4M_rep2 |
| CCGTTCACATCGATAC-1 | WT_4M_rep2 |
| CCGTTCATCTGAATGC-1 | WT_4M_rep2 |
| CCTATCGTCCGATAAC-1 | WT_4M_rep2 |
| CCTATCGTCCGCATAA-1 | WT_4M_rep2 |
| CCTCACAAGCCGTCGT-1 | WT_4M_rep2 |
| CCTCACATCATAAGGA-1 | WT_4M_rep2 |
| CCTCATGTCAATGTCG-1 | WT_4M_rep2 |
| CCTGCATTCTGATGGT-1 | WT_4M_rep2 |
| CGAAGTTAGCGACATG-1 | WT_4M_rep2 |
| CGAAGTTCAGACTCTA-1 | WT_4M_rep2 |
| CGGACACAGCACACAG-1 | WT_4M_rep2 |
| CGGGCATCACCTGTCT-1 | WT_4M_rep2 |
| CGTGCTTGTTCAAACC-1 | WT_4M_rep2 |
| CTAAGTGTCGAGGCAA-1 | WT_4M_rep2 |

| | |
|---|---|
| CTACATTTCCTAGCTC-1 | WT_4M_rep2 |
| CTACGGGGTTTACTGG-1 | WT_4M_rep2 |
| CTACGGGTCGGACGTC-1 | WT_4M_rep2 |
| CTCATTACATTGCCTC-1 | WT_4M_rep2 |
| CTCCAACTCGCTGATA-1 | WT_4M_rep2 |
| CTCTGGTCACAAATAG-1 | WT_4M_rep2 |
| CTGTACCGTGATTCTG-1 | WT_4M_rep2 |
| CTTCCTTGTATACCCA-1 | WT_4M_rep2 |
| GAAACCTTCTAGAACC-1 | WT_4M_rep2 |
| GAAGAATTCTCCCAAC-1 | WT_4M_rep2 |
| GAAGCCCGTTGTCCCT-1 | WT_4M_rep2 |
| GAAGCGAGTAGGAGTC-1 | WT_4M_rep2 |
| GAATAGATCTGCCTGT-1 | WT_4M_rep2 |
| GAATCGTCAAACTAGA-1 | WT_4M_rep2 |
| GACCTTCTCTCATGGA-1 | WT_4M_rep2 |
| GACGTTATCCTGTTGC-1 | WT_4M_rep2 |
| GACTTCCAGAGAACCC-1 | WT_4M_rep2 |
| GAGGGATGTAGATCGG-1 | WT_4M_rep2 |
| GAGGGTAGTTATTCCT-1 | WT_4M_rep2 |
| GAGTGTTCAAATAAGC-1 | WT_4M_rep2 |
| GATAGCTGTCGTTGCG-1 | WT_4M_rep2 |
| GATCGTAAGGTAGTAT-1 | WT_4M_rep2 |
| GCACATAGTACGACAG-1 | WT_4M_rep2 |
| GCACGGTTCTTTCGAT-1 | WT_4M_rep2 |
| GCACTAACAATATCCG-1 | WT_4M_rep2 |
| GCACTAACACGCGCAT-1 | WT_4M_rep2 |
| GCATGATGTCAAATCC-1 | WT_4M_rep2 |
| GCGGAAATCCCATAAG-1 | WT_4M_rep2 |
| GCTCAAATCGACCAAT-1 | WT_4M_rep2 |
| GCTGCAGAGCGAGAAA-1 | WT_4M_rep2 |
| GCTTTCGTCGCCGAGT-1 | WT_4M_rep2 |
| GGAACCCGTTTACCAG-1 | WT_4M_rep2 |
| GGAGAACGTAATGTGA-1 | WT_4M_rep2 |
| GGAGCAACAATACGCT-1 | WT_4M_rep2 |
| GGAGGATCATCTCATT-1 | WT_4M_rep2 |
| GGCAGTCGTTCCGCAG-1 | WT_4M_rep2 |
| GGGACTCGTGGCAACA-1 | WT_4M_rep2 |
| GGGCTCAAGGCATCTT-1 | WT_4M_rep2 |
| GGGTCACTCAGGAAAT-1 | WT_4M_rep2 |
| GGGTGAATCGGATTAC-1 | WT_4M_rep2 |
| GGTGGCTGTTGCATAC-1 | WT_4M_rep2 |
| GGTGTCGTCCATACAG-1 | WT_4M_rep2 |
| GTAATCGTCTCATTGT-1 | WT_4M_rep2 |
| GTACAGTCAGTGTGGA-1 | WT_4M_rep2 |

| | |
|---|---|
| GTATTTCCAGTATGAA-1 | WT_4M_rep2 |
| GTATTTCTCGTCGATA-1 | WT_4M_rep2 |
| GTCAGCGAGGTTCACT-1 | WT_4M_rep2 |
| GTCATCCCAAATGAAC-1 | WT_4M_rep2 |
| GTCATTTGTGAGTTGG-1 | WT_4M_rep2 |
| GTCGTAACAGTAACAA-1 | WT_4M_rep2 |
| GTGCGTGGTAGTGGCA-1 | WT_4M_rep2 |
| GTGCTTCCATATAGCC-1 | WT_4M_rep2 |
| GTGGAAGTCACGGACC-1 | WT_4M_rep2 |
| GTGGTTAGTCACAGAG-1 | WT_4M_rep2 |
| GTGTCCTTCCCGAATA-1 | WT_4M_rep2 |
| GTGTGGCAGCAAGTGC-1 | WT_4M_rep2 |
| GTTAGACGTAGCGTCC-1 | WT_4M_rep2 |
| GTTCGCTCAACGATCT-1 | WT_4M_rep2 |
| GTTGCTCTCAATCTCT-1 | WT_4M_rep2 |
| GTTGTCCTCGGCTGTG-1 | WT_4M_rep2 |
| GTTTACTCAACAGTGG-1 | WT_4M_rep2 |
| TAACGACAGCTGTTCA-1 | WT_4M_rep2 |
| TAAGTCGAGAATTGCA-1 | WT_4M_rep2 |
| TAATCTCGTGTCCAAT-1 | WT_4M_rep2 |
| TACCTCGGTGCTATTG-1 | WT_4M_rep2 |
| TACGCTCCAACTGTGT-1 | WT_4M_rep2 |
| TAGAGTCAGCACGTCC-1 | WT_4M_rep2 |
| TAGATCGCATTGCTGA-1 | WT_4M_rep2 |
| TAGCACATCTTTCAGT-1 | WT_4M_rep2 |
| TAGGAGGTCCTCTCTT-1 | WT_4M_rep2 |
| TAGGTACGTACGGGAT-1 | WT_4M_rep2 |
| TAGGTTGCAGATACCT-1 | WT_4M_rep2 |
| TATACCTAGAAGGCTC-1 | WT_4M_rep2 |
| TATACCTTCTTGGCTC-1 | WT_4M_rep2 |
| TATATCCAGTTGCCTA-1 | WT_4M_rep2 |
| TATCCTAGTAAGCGGT-1 | WT_4M_rep2 |
| TATCGCCAGACAGTCG-1 | WT_4M_rep2 |
| TATCTTGGTAGCTTGT-1 | WT_4M_rep2 |
| TATGTTCGTGGCCACT-1 | WT_4M_rep2 |
| TCAAGCAGTTCCTAAG-1 | WT_4M_rep2 |
| TCAAGTGGTGCCTAAT-1 | WT_4M_rep2 |
| TCAGGGCCATGGAATA-1 | WT_4M_rep2 |
| TCAGTTTAGGTCGAGT-1 | WT_4M_rep2 |
| TCATCCGCACAAAGCG-1 | WT_4M_rep2 |
| TCATGAGGTTTGAAAG-1 | WT_4M_rep2 |
| TCATGTTGTTGAGGAC-1 | WT_4M_rep2 |
| TCATTTGTCCATGCAA-1 | WT_4M_rep2 |
| TCCACGTTCGGACCAC-1 | WT_4M_rep2 |

| | |
|---|---|
| TCCATGCTCACGGGCT-1 | WT_4M_rep2 |
| TCCGAAAAGGTTTGAA-1 | WT_4M_rep2 |
| TCCTCTTGTACGATTC-1 | WT_4M_rep2 |
| TCCTGCAGTGGCAGAT-1 | WT_4M_rep2 |
| TCGAAGTAGCACACCC-1 | WT_4M_rep2 |
| TCGACCTTCCAAAGGG-1 | WT_4M_rep2 |
| TCGGATAAGTAAGACT-1 | WT_4M_rep2 |
| TCGGATAGTCCATCTC-1 | WT_4M_rep2 |
| TCGGGCATCTACAGGT-1 | WT_4M_rep2 |
| TCGGTCTAGACCCGCT-1 | WT_4M_rep2 |
| TCGGTCTGTACCCACG-1 | WT_4M_rep2 |
| TCGTGGGGTAGGACCA-1 | WT_4M_rep2 |
| TCTCAGCAGCAACAGC-1 | WT_4M_rep2 |
| TCTCAGCGTCTTTATC-1 | WT_4M_rep2 |
| TCTTGCGTCAAATAGG-1 | WT_4M_rep2 |
| TGACAGTCAGGTTCCG-1 | WT_4M_rep2 |
| TGATCAGAGAGTGTGC-1 | WT_4M_rep2 |
| TGCACGGCAGAAGCTG-1 | WT_4M_rep2 |
| TGCAGGCAGGAACGTC-1 | WT_4M_rep2 |
| TGCGGCATCCCAAGCG-1 | WT_4M_rep2 |
| TGCGGGTCACCGGCTA-1 | WT_4M_rep2 |
| TGGTAGTCATGGAGAC-1 | WT_4M_rep2 |
| TGTAAGCAGTCCCAAT-1 | WT_4M_rep2 |
| TGTAGACAGATACTGA-1 | WT_4M_rep2 |
| TGTTCATCACCCAATA-1 | WT_4M_rep2 |
| TGTTGGATCCGCAACG-1 | WT_4M_rep2 |
| TTACTGTAGTTGGCTT-1 | WT_4M_rep2 |
| TTAGGCAAGACCATTC-1 | WT_4M_rep2 |
| TTAGGGTCAAGCACAG-1 | WT_4M_rep2 |
| TTCCGTGAGTTGGACG-1 | WT_4M_rep2 |
| TTCCTTCCAGCGAACA-1 | WT_4M_rep2 |
| TTCGCTGCACATGACT-1 | WT_4M_rep2 |
| TTCTAGTGTCACTCTC-1 | WT_4M_rep2 |
| TTCTTCCTCCGAACGC-1 | WT_4M_rep2 |
| TTGAACGAGAAGCCAC-1 | WT_4M_rep2 |
| TTGACCCCACTAGAGG-1 | WT_4M_rep2 |
| TTGCATTGTCTTTCAT-1 | WT_4M_rep2 |
| TTGCCTGTCAAGCGTT-1 | WT_4M_rep2 |
| TTGTGGATCTCACCCA-1 | WT_4M_rep2 |
| TTTACGTTCCTGTTGC-1 | WT_4M_rep2 |
| TTTAGTCGTCGGATTT-1 | WT_4M_rep2 |
| TTTCATGAGCCTATCA-1 | WT_4M_rep2 |
| TTTGATCCAGGCACTC-1 | WT_4M_rep2 |

Table S1. The table shows the cell barcodes of the cone photoreceptors.
